# No substantial pre-existing B cell immunity against SARS-CoV-2 in healthy adults

**DOI:** 10.1101/2021.09.08.459398

**Authors:** Meryem Seda Ercanoglu, Lutz Gieselmann, Sabrina Dähling, Nareshkumar Poopalasingam, Susanne Detmer, Manuel Koch, Michael Korenkov, Sandro Halwe, Michael Klüver, Veronica Di Cristanziano, Hanna Janicki, Maike Schlotz, Johanna Worczinski, Birgit Gathof, Henning Gruell, Matthias Zehner, Stephan Becker, Kanika Vanshylla, Christoph Kreer, Florian Klein

**Affiliations:** Laboratory of Experimental Immunology, Institute of Virology, Faculty of Medicine and University Hospital Cologne, University of Cologne, 50931 Cologne, Germany; German Center for Infection Research, Partner Site Bonn-Cologne, 50931 Cologne, Germany; Center for Molecular Medicine Cologne (CMMC), University of Cologne, 50931 Cologne, Germany; Institute for Dental Research and Oral Musculoskeletal Biology and Center for Biochemistry, University of Cologne, 50931 Cologne, Germany; Institute of Transfusion Medicine, Faculty of Medicine and University Hospital Cologne, 50937 Cologne, Germany; Institute of Virology, Philipps University Marburg, Hans-Meerwein-Straße 2, 35042 Marburg, Germany; German Center for Infection Research, Partner Site Giessen-Marburg-Langen, 35043 Marburg, Germany

## Abstract

Pre-existing immunity against SARS-CoV-2 may have critical implications for our understanding of COVID-19 susceptibility and severity. Various studies recently provided evidence of pre-existing T cell immunity against SARS-CoV-2 in unexposed individuals. In contrast, the presence and clinical relevance of a pre-existing B cell immunity remains to be fully elucidated. Here, we provide a detailed analysis of the B cell response to SARS-CoV-2 in unexposed individuals. To this end, we extensively investigated the memory B cell response to SARS-CoV-2 in 150 adults sampled pre-pandemically. Comprehensive screening of donor plasma and purified IgG samples for binding and neutralization in various functional assays revealed no substantial activity against SARS-CoV-2 but broad reactivity to endemic betacoronaviruses. Moreover, we analyzed antibody sequences of 8,174 putatively SARS-CoV-2-reactive B cells on a single cell level and generated and tested 158 monoclonal antibodies. None of the isolated antibodies displayed relevant binding or neutralizing activity against SARS-CoV-2. Taken together, our results show no evidence of relevant pre-existing antibody and B cell immunity against SARS-CoV-2 in unexposed adults.

## Introduction

The current pandemic of severe acute respiratory syndrome coronavirus 2 (SARS-CoV-2) represents a global health emergency that challenges health care systems throughout the world. SARS-CoV-2 infections appear with a broad spectrum of clinical manifestations ranging from asymptomatic infections to life-threatening acute respiratory distress syndrome (ARDS), multi organ failure, septic shock, and death. Although influenced by multiple contributing factors, disease severity is substantially shaped by innate and adaptive immune responses. Pre-existing immunity against SARS-CoV-2 could represent a crucial determinant of disease severity and clinical outcome. For example, recognition of SARS-CoV-2 by a pre-existing background immunity could limit disease severity by rapidly mounting specific immune responses. However, pre-existing immunity may also be detrimental for the clinical course by mechanisms of antibody-dependent enhancement (ADE) (Khurana et al., 2013; de Alwis et al., 2014; Katzelnick et al., 2017; Arvin et al., 2020) or original antigenic sin (OAS) (Vatti et al., 2017), which have been previously described for other viral pathogens, such as dengue (de Alwis et al., 2014; Katzelnick et al., 2017; Mongkolsapaya et al., 2003; Midgley et al., 2011; Rothman, 2011) and influenza viruses (Linderman and Hensley, 2016; Zhang et al., 2019; Arevalo et al., 2020).

Pre-existing T cell immune responses against SARS-CoV-2 have been observed in unexposed individuals (Mateus et al., 2020; Grifoni et al., 2020; Le Bert et al., 2020; Braun et al., 2020; Bacher et al., 2020; Weiskopf et al., 2020; Echeverría et al., 2021). In these studies, T cell reactivity against the spike (S) and nucleocapsid (N) proteins as well the non-structural proteins NSP7 and NSP13 was determined using antigen peptide pools (Grifoni et al., 2020; Le Bert et al., 2020; Mateus et al., 2020). Importantly, pronounced T cell reactivity was detected against S protein peptides exhibiting a high degree of homology to endemic ‘common cold’ human coronaviruses (HCoV), including HCoV-OC43, HCoV-HKU-1, HCoV-NL63, and HCoV-229E. Therefore, pre-existing T cell immunity against SARS-CoV-2 is hypothesized to originate from prior exposure to endemic HCoVs (Mateus et al., 2020; Grifoni et al., 2020; Le Bert et al., 2020; Braun et al., 2020; Weiskopf et al., 2020).

Pre-existing B cell immunity may be already germline-encoded in the naïve B cell repertoire or originate from cross-reactive immune responses against related pathogens or variants. Many potent SARS-CoV-2 neutralizing antibodies (nAbs) exhibit binding by germline-encoded amino acid residues within the complementarity-determining regions 1 and 2 (CDRH1 and CDRH2) (Barnes et al., 2020a; Hurlburt et al., 2020; Shi et al., 2020; Wu et al., 2020; Yuan et al., 2020), are restricted to specific heavy chain V genes (Brouwer et al., 2020; Cao et al., 2020; Ju et al., 2020; Robbiani et al., 2020; Rogers et al., 2020; Seydoux et al., 2020; Wu et al., 2020; Zost et al., 2020), or exhibit a low degree of somatic mutations (Barnes et al., 2020b; Kreer et al., 2020a; Robbiani et al., 2020; Seydoux et al., 2020). This suggests that near-germline B cell receptor (BCR) sequences with close similarity to SARS-CoV-2 reactive antibodies are already encoded in the naïve B cell repertoire and can be readily selected to mount a potent B cell response without further affinity maturation. In line with this, we previously identified potential heavy and/or light-chain precursor sequences of SARS-CoV-2-binding as well as -neutralizing antibodies by deep sequencing of naïve B cell receptor repertoires sampled before the SARS-CoV-2 pandemic (Kreer et al., 2020a).

Beyond germline encoded immunity, cross-reactive immune responses to endemic HCoVs may also account for a pre-existing humoral or B cell immunity against SARS-CoV-2. Recent studies provide controversial results regarding the frequency of cross-reactive antibodies in the sera of unexposed individuals (Anderson et al., 2021; Ng et al., 2020; Nguyen-Contant et al., 2020; Poston et al., 2020; Shrock et al., 2020; Song et al., 2021) and their association with protection from disease severity or hospitalization (Anderson et al., 2021; Gombar et al., 2021; Sagar et al., 2021). Whereas one study found SARS-CoV-2 reactive antibodies in a considerable number of unexposed individuals -particularly among children, adolescents and pregnant women (Ng et al., 2020)– other studies did not find comparable evidence (Anderson et al., 2021; Nguyen-Contant et al., 2020; Poston et al., 2020; Song et al., 2021). All studies to date depended on the investigation of SARS-CoV-2 reactivity in serum/plasma or affinity-enriched or secreted IgG fractions of unexposed individuals (Anderson et al., 2021; Ng et al., 2020; Nguyen-Contant et al., 2020; Poston et al., 2020; Shrock et al., 2020; Song et al., 2021). These conflicting results call for more comprehensive and detailed investigations which beyond plasma and IgG fractions also involve BCR sequence analysis of SARS-CoV-2 reactive B cells and characterization of recombinant monoclonal antibodies.

In order to investigate the presence of a relevant pre-existing SARS-CoV-2 B cell immunity, we extensively investigated plasma samples, single B cells, and monoclonal antibodies isolated from 150 SARS-CoV-2 unexposed individuals. We found no evidence of a pre-existing B cell immunity that may account for the broad clinical spectrum of SARS-CoV-2 infections. Instead, our results indicate that rare naïve B cell precursors are selected to mount a SARS-CoV-2 directed antibody response upon infection.

## Results

### Pre-pandemic naïve B cell receptor repertoires encode for heavy and light chains of SARS-CoV-2 reactive antibodies

By performing NGS analyses on healthy individuals, we previously identified rare heavy and light chain variable regions of pre-pandemic naive B cells that closely resembled near-germline SARS-CoV-2-reactive antibodies (Kreer et al., 2020a). This raised the question, whether naïve B cells can encode for antibodies that do not require further affinity maturation for a potent SARS-CoV-2 reactivity. To test this hypothesis, we cloned and expressed 31 pre-pandemic heavy or light chains obtained from naïve B cells together with heavy or light chains from SARS-CoV-2-reactive mAbs derived from convalescent individuals and tested these chimeras for S protein binding and SARS-CoV-2 neutralization activity (**Figure 1**). V gene and CDR3 differences between NGS-derived and original chains ranged from 0 to 5 and 0 to 3 amino acids, respectively (**Table S2**). Except for mAbs CnC2t1p1_B4 and MnC2t1p1_C12, pairing of pre-pandemic light chains with the original heavy chains did retain binding and/or neutralization activity. For 20 out of 23 chimeric mAbs, pairing of pre-pandemic heavy chains with the original light chain led to a loss of binding and neutralization activity. However, for one antibody (MnC2t1p1_C12) pairing of the original light chain with 3 out of 13 pre-pandemic heavy chains was able to retain binding activity suggesting that these heavy chains can assemble SARS-CoV-2 spike protein reactive antibodies. We conclude that some naïve B cells can express heavy or light chains that can be components of SARS-CoV-2-reactive antibodies without the need for further affinity maturation.

**Figure 1:**
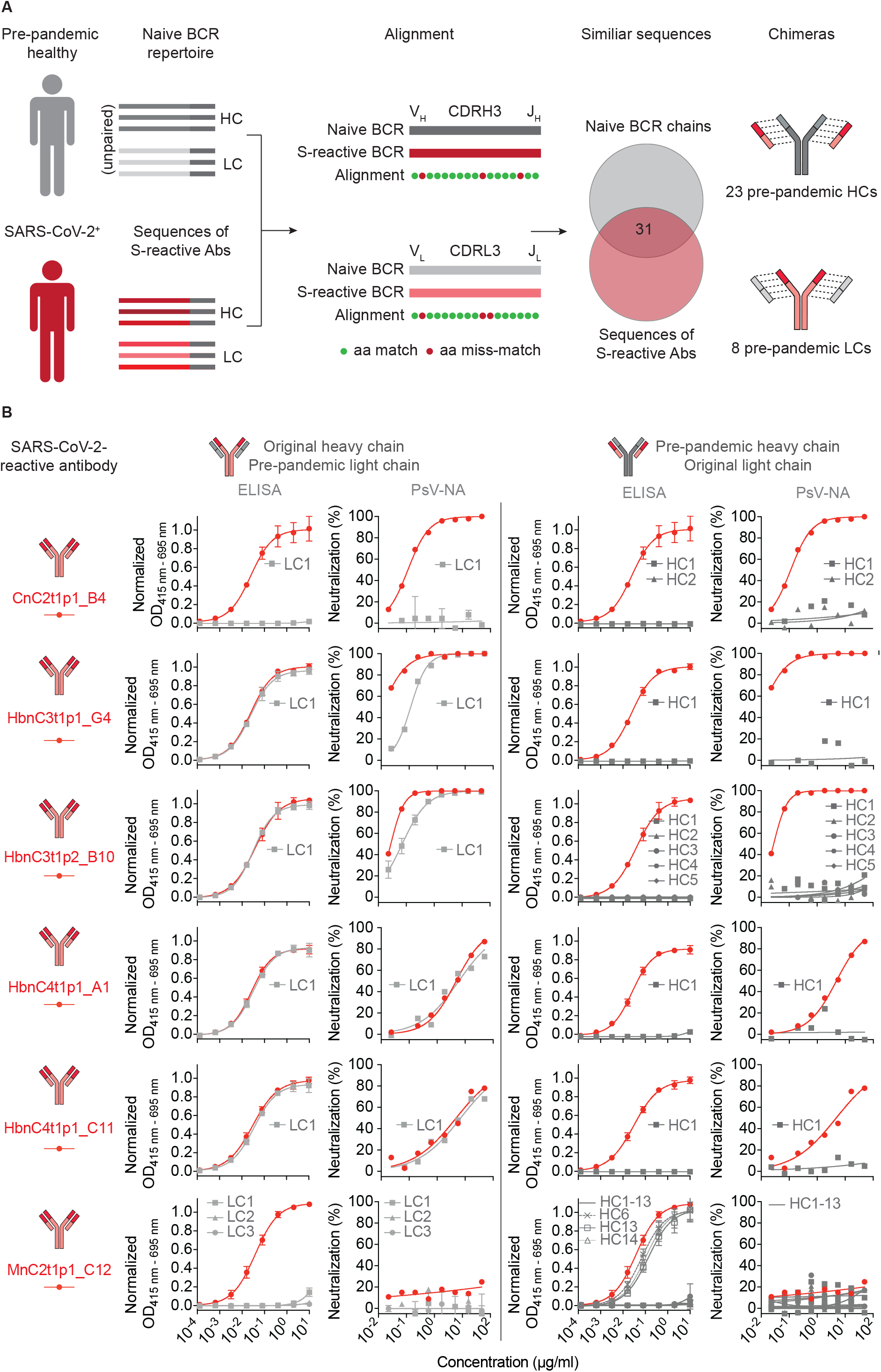
Pre-pandemic naïve B cells express SARS-CoV-2-reactive heavy or light chains. **(A**) Selection of heavy and light chain sequences in a pre-pandemic naïve B cell repertoire that closely resemble heavy and light chains of SARS-CoV-2 reactive antibodies (Abs). **(B)** Binding and neutralization plots against SARS-CoV-2 of 6 previously described SARS-CoV-2-reactive antibodies (red) and 31 chimeric antibodies composed of heavy or light chain of the original SARS-CoV-2-reactive antibodies paired with a NGS-derived heavy (darker grey) or light (grey) chain. Activity of mAbs against SARS-CoV-2 was determined by serial dilution ELISA and pseudotyped neutralization assays (PsV-NA). Each antibody was produced in at least duplicates. ELISAs and neutralization assays were performed with biological replicates. Symbols depict means and error bars indicate standard deviation.

### Pre-pandemic plasma samples and polyclonal IgG exhibit no significant reactivity to SARS-CoV-2

To investigate whether SARS-CoV-2-reactive antibodies are present in the plasma of unexposed individuals, we investigated pre-pandemic blood samples from 150 donors. Samples were collected between August and November 2019 and were studied for binding and neutralization activity against SARS-CoV-2 (**Figure 2A** and **B**; **Table S1**). Donors were between 18 and 66 years old (with a mean/median age of 30.6/27 years) (**Figure 2A** and **Table S1**). 49.3 % of donors were male and 50.7 % female (**Figure 2A** and **Table S1**). Plasma samples of all 150 donors were tested for binding to the soluble full trimeric SARS-CoV-2 S ectodomain (S1/S2) or the S1 subunit (S1) by commercially available (com. IA) and in-house (ELISA) immunoassays. (**Figures 2B, C** and **Figure S1**). In addition, binding activity was assessed against cell surface expressed full-length SARS-CoV-2 S protein by flow cytometry (cell assay, CA; **Figures 2B** and **C**). Plasma samples showed mostly no or only minimal binding of IgG, IgM and IgA to soluble or cell surface expressed SARS-CoV-2 S proteins (**Figures 2C** and F**igure S1**). Only in a few samples (Pre033 IgA, Pre051 IgG, Pre074 IgA, Pre084 IgM and Pre097 IgA) notable binding was detected in one single assay (**Figure 2C** and **Figure S2**). However, binding detected by immunoassays could not be confirmed by flow-cytometry against cell surface expressed full-length S protein (**Figure 2C**). In CAs, one donor (Pre004) exhibited weak plasma IgG activity to full lengths S protein. Furthermore, 1:10 dilutions of 150 plasma samples were screened for neutralization activity against SARS-CoV-2 wildtype-(WT) and pseudovirus (PSV) (**Figures 2C** and **Figure S2**). Samples from two donors neutralized SARS-CoV-2 pseudovirus by 50 – 60 %. However, neutralization activity of these samples could not be confirmed by testing of serial dilutions or wildtype neutralization assays (**Figure 2C** and **Figure S2**).

**Figure 2:**
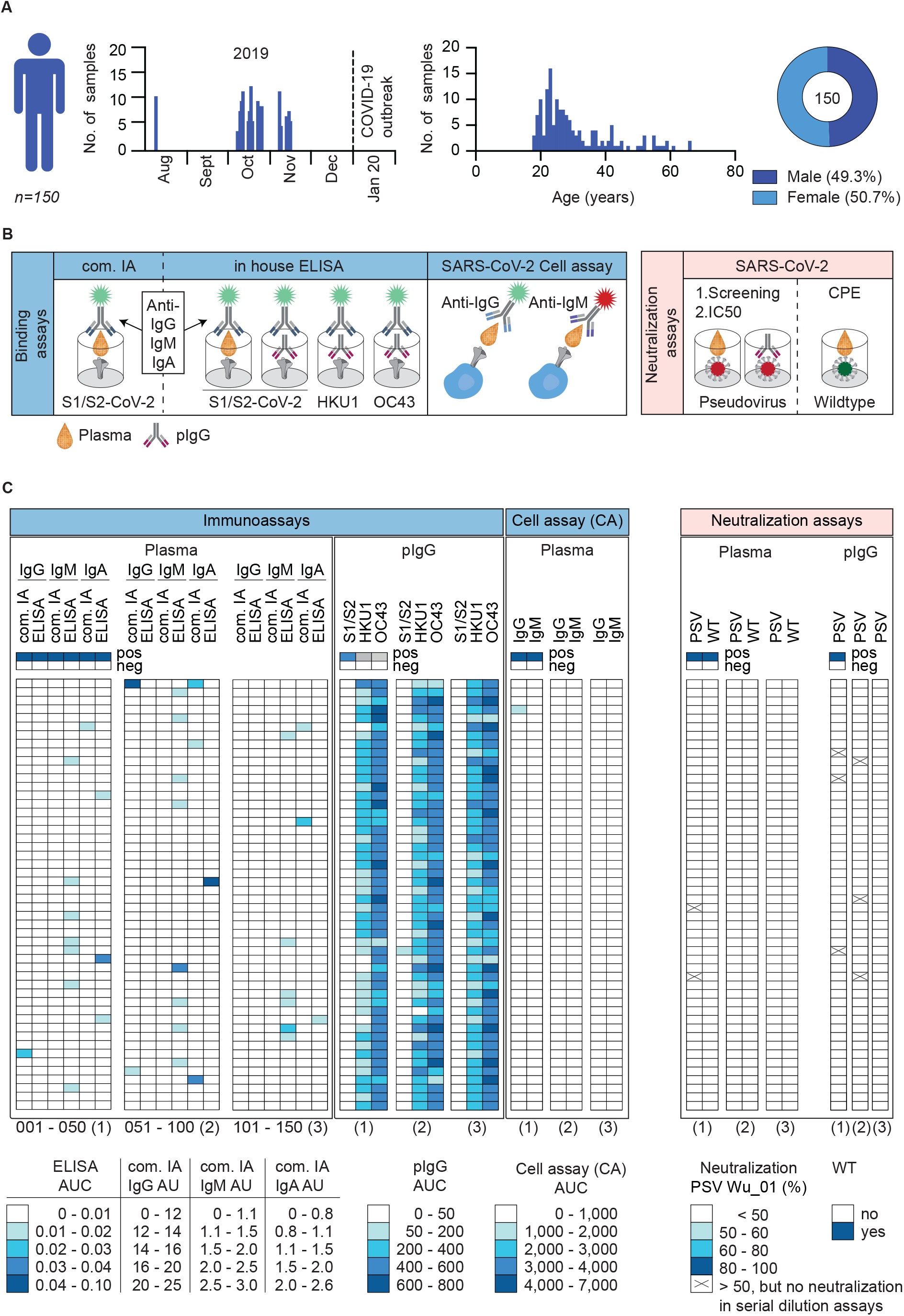
Screening of pre-pandemic samples from 150 adults reveal no relevant reactivity against SARS-CoV-2. **(A)** Timeline of blood collections and demographic characteristics of 150 donors sampled before the SARS-CoV-2 pandemic. Pre-pandemic blood samples were collected as buffy coats between August and November of 2019. Gender and age distribution of donors are illustrated as pie chart and bar plots. **(B)** Pre-pandemic plasma samples and purified IgGs (pIgG) were tested for binding and neutralization using different experimental approaches. Plasma IgG, IgM and IgA as well as pIgG were tested for binding to SARS-CoV-2, HKU1 and OC43 S proteins using in house (ELISA) and commercially available immunoassays (com. IA). Binding of plasma IgG and IgM to cell surface expressed SARS-CoV-2 S protein was also determined by FACS analysis (CA). Neutralization activity was determined against SARS-CoV-2 pseudo- and wildtype virus (PSV or WT). **(C)** Heat map visualizing binding (AUC or AU) and neutralization activity (% or CPE) of pre-pandemic plasma samples and pIgG against SARS-CoV-2 and endemic HCoVs (HKU1 and OC43) S proteins or SARS-CoV-2 wildtype (WT) and pseudovirus (PSV Wu_01), respectively (see also **Figure S1 and S2**). Immunoassays were performed in duplicate experiments and the AUC is presented as geometric mean of duplicates. Neutralization activity was first determined for single sample concentrations. Samples that displayed neutralization activity of ≥ 50% in single concentrations were repeatedly investigated in serial dilutions (x) (see also **Figure S2**). Samples were tested in duplicates in single experiments. The average of neutralization is presented and each row represents one donor.

Next, polyclonal IgGs (pIgG) were purified from pre-pandemic plasma samples and tested for binding to soluble full trimeric S proteins of SARS-CoV-2 and endemic human-pathogenic betacoronaviruses (HKU1 and OC43) (**Figures 2C** and **Figure S2**). Although pIgG showed strong reactivity to HKU1 and OC43 S proteins, no binding to SARS-CoV-2 trimeric S protein was detected. Furthermore, neutralization activity of pIgG against SARS-CoV-2 pseudovirus was determined at a concentration of 1 mg/ml. PIgG of six donors exhibited neutralization activity of SARS-CoV-2 pseudovirus by ≥ 50 %. However, neutralization activity of these samples could not be confirmed by testing serial dilutions (**Figure 2C** and **Figure S2**). We conclude that pre-pandemic plasma and polyclonal IgG samples obtained from 150 adults before the onset of the pandemic exhibit only minimal reactivity to SARS-CoV-2 in few samples and overall no SARS-CoV-2-neutralizing activity.

### No detection of SARS-CoV-2-reactive B cells in pre-pandemic samples

The lack of binding or neutralization activity against SARS-CoV-2 on plasma level does not exclude the existence of SARS-CoV-2-reactive B cells in unexposed individuals. To investigate the presence of SARS-CoV-2 specific B cells, we performed single B cell sorts of 40 donors sampled before the pandemic (**Figure 3** and **Figure S3**). Using the same analysis gate as for COVID-19 convalescent donors, frequencies of SARS-CoV-2-reactive IgG^+^ and IgG^−^ B cells isolated from pre-pandemic blood samples were significantly lower (p value < 0.0001). For COVID-19 convalescent donors, frequencies ranged from 0.002 to 0.065 % for IgG^+^ (median 0.02 %) and 0.007 to 0.39 % for IgG^−^ (median 0.031 %) B cells (**Figure 3A**). Applying the same analysis gate of the COVID-19 convalescent samples to our pre-pandemic samples, frequencies ranged from 0 to 0.0013 % for IgG^+^ (median 0.0001 %) and from 0 to 0.016 for IgG^−^ (median 0.003 %) B cells (**Figure 3A** and **Figure S3**). Therefore, we conclude that, if present at all, SARS-CoV-2-reactive B cells have a significantly lower frequency in pre-pandemic samples. We reasoned that gate settings applied for COVID-19 convalescent samples may exclude reactive B cells with low spike affinity which may be present in individuals unexposed to SARS-CoV-2. To assure that such cells do not remain undetected, we adjusted the actual sorting gate (**Figure 3A**) to isolate a total of 8,174 putatively SARS-CoV-2-reactive single B cells, of which 3,852 were IgG^+^ and 4,322 IgG^−^ cells. Of those we amplified a total of 5,432 productive heavy chain sequences, of which 2,789 sequences accounted for IgG and 2,643 for IgM heavy chains, respectively (**Figure 3B**). Sequence analyses showed that in each individual 0 to 58 % of the sequences were clonally related with a mean clone number of 8.1 clones per individual and a mean clone size of 2.9 members per clone (**Figure 3B**). Heavy chain variable (V_H_) gene segment distribution, heavy chain complementarity-determining region 3 (CDRH3) length and V_H_ gene germline identities showed no notable divergence to a reference memory IgG and naïve IgM repertoire data set (Kreer et al., 2020a) (**Figure 3C**). The lack of this divergence in combination with the FACS data suggest the absence of SARS-CoV-2-reactive B cells in pre-pandemic samples.

**Figure 3:**
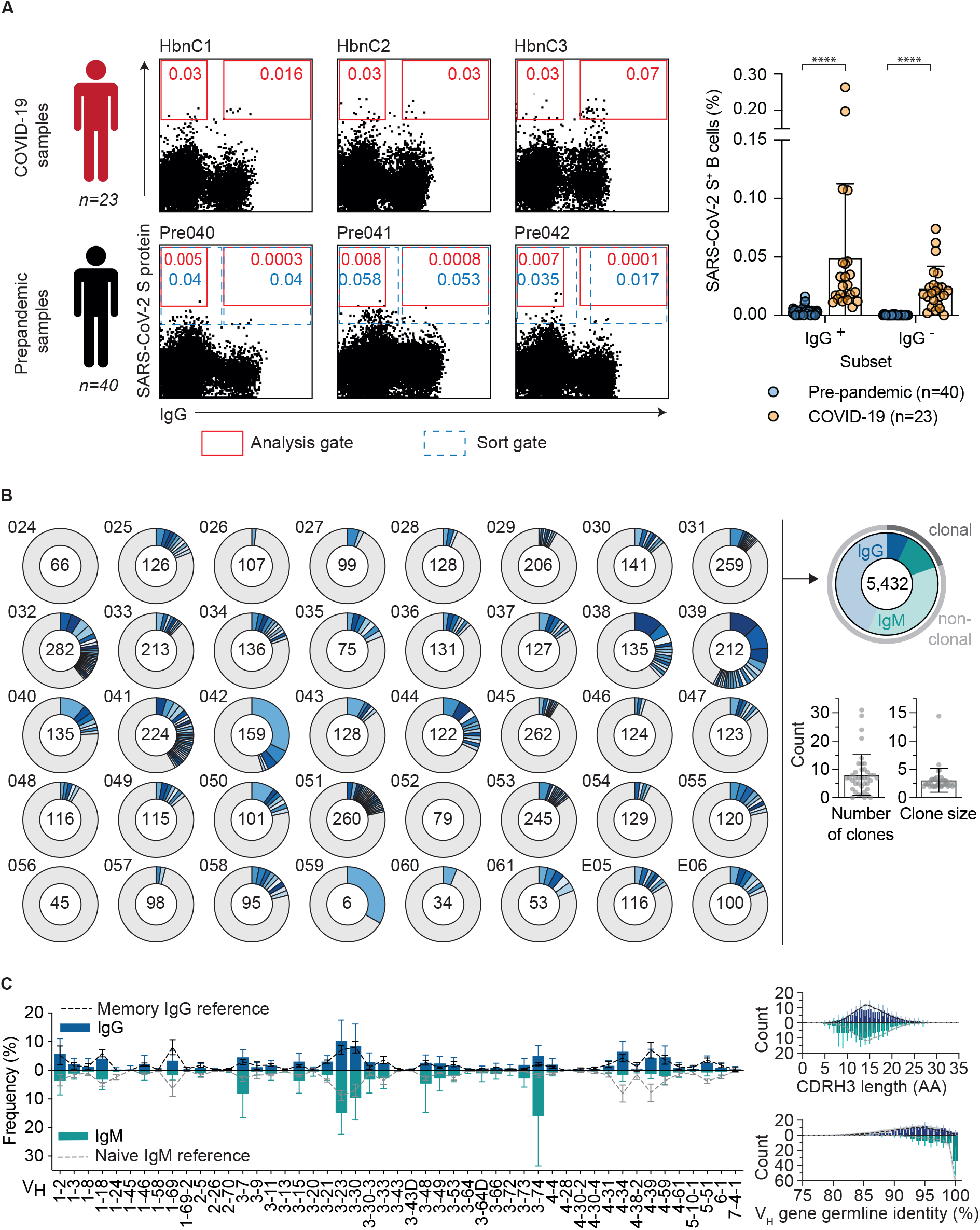
Lack of SARS-CoV-2-reactive B cells and distinct antibody sequence features in pre-pandemic samples. **(A)** Representative dot plots of SARS-CoV-2-reactive, CD19^+^CD20^+^, IgG^+^ and IgG^−^ B cells of COVID-19 samples compared to pre-pandemic samples. Depicted numbers indicate frequencies of S protein reactive B cells (see also **Figure S3**). Red colored gate indicates gating strategy for analysis and doted gate indicates actual sorting gate. Dot plot bar graph displays the mean ± SD frequency of SARS-CoV-2-reactive, IgG^+^ and IgG^−^ B cells in 40 pre-pandemic and 23 COVID-19 samples (*P < 0.05, **P < 0.01, ***P < 0.001 and ****P < 0.0001; unpaired two-tailed t test). **(B)** Clonal relationship of heavy chain sequences amplified from single SARS-CoV-2-reactive IgG^+^ and IgG^−^ B cells isolated from 40 donors sampled before the pandemic. Individual clones are colored in shades of blue, grey and white. In the center of each pie chart, numbers of productive heavy chain sequences are illustrated. Presentation of clone sizes are proportional to the total number of productive heavy chain sequences per clone. **(C)** V_H_ gene distribution, V_H_ gene germline identity and CDRH3 length distribution in amino acids (AA) were separately determined for IgG and IgM. Distributions were calculated per individual. Bar and line plots show mean ± SD.

### Monoclonal antibodies isolated from pre-pandemic samples are not reactive against SARS-CoV-2

To confirm the absence of SARS-CoV-2-reactive B cells in pre-pandemic samples on a functional level, we selected 200 antibody candidates among 36 donors from single cell sorted B cells for production and functional testing (**Figure 4A**). Selection was conducted based on sequence similarity (see **Methods section**) to 920 SARS-CoV-2-reactive antibodies deposited in the CoVAbDab (n = 18) and on random sequence selection (n = 182) (**Figure 4**). Criteria for the similarity selection were identical VH/JH combinations, low level of CDRH3 length differences, and Levenshtein distances between isolated and deposited BCR sequences. The random selection included clonal as well as non-clonal sequences and was performed to ensure an equal selection of BCR sequences among different donors and clonotypes. In total, we successfully produced 158 monoclonal antibodies (81 IgM-, 77 IgG-derived) as IgG1 isotypes for functional testing. First, we determined binding activity of these antibodies to SARS-CoV-2 S protein and evaluated cross-reactivity to HKU1 and OC43 S proteins by ELISA (**Figure 4B**). Monoclonal antibodies showed no relevant binding or cross-reactivity against SARS-CoV-2, HKU-1 or OC43 S proteins. Next, we tested all 158 monoclonal antibodies for neutralization activity against SARS-CoV-2 pseudovirus in single concentrations of 50 μg/ml (**Figure S4**) and in serial dilutions (**Figure 4B).** In line with the lack of binding activity, none of the produced antibodies showed neutralizing activity up to concentrations of 50 μg/ml (**Figure 4B** and **Figure S4**). We conclude that putatively SARS-CoV-2-S^+^ B cells from pre-pandemic samples do not encode for SARS-CoV-2-reactive B cell receptors.

**Figure 4:**
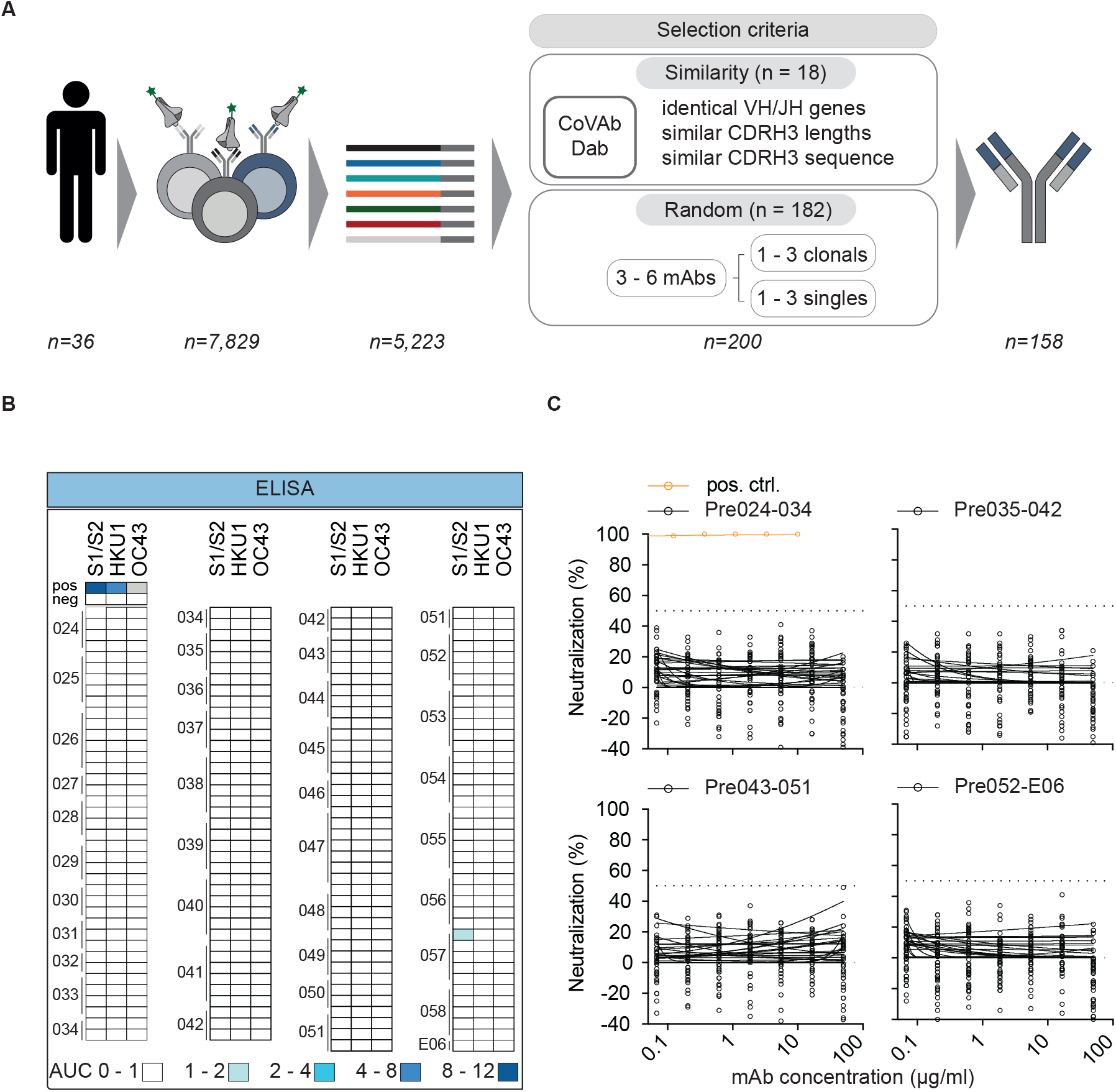
Monoclonal antibodies isolated from pre-pandemic samples are not reactive to SARS-CoV-2 and endemic HCoVs. **(A)** Illustration demonstrating the sequence selection for downstream antibody production. Sequences were selected for antibody production based on similarity to antibody sequences deposited in the CoVAbDab and on random selection. From 8,174 SARS-CoV-2-reactive IgG+ and IgG-B cells, 5,432 productive heavy chain sequences were amplified. For antibody production, 18 HC sequences were selected based on similarity selection and 182 HC sequences were selected based on random selection. In total 158 antibodies were produced. **(B)** Heat map visualizing binding (AUC) and neutralization activity (%) of monoclonal antibodies isolated from pre-pandemic blood samples against SARS-CoV-2 and endemic HCoVs (HKU1 and OC43) S proteins or SARS-CoV-2 pseudovirus (PSV Wu_01), respectively (see also **Figure S4**). Each row represents one monoclonal antibody. ELISAs were performed in duplicate experiments and the AUC is presented as geometric mean of duplicates. Neutralization activity was determined for single sample concentrations. Samples were tested in duplicates and the average of neutralization is presented. **(C)** Neutralization activity against SARS-CoV-2 pseudovirus (PSV Wu_01) was verified for all mAbs in serial dilutions.

## Discussion

Adaptive immune responses against a pathogen are shaped by the naïve immune repertoire and imprinted by previous encounters to the same pathogen or a related variant. Investigation of pre-existing immunity to SARS-CoV-2 can advance our understanding of protective immunity, susceptibility to infection, disease severity and guide the development of vaccination strategies (Imkeller and Wardemann, 2018; Niu et al., 2020; Schultheiß et al., 2020). Although various studies already provide evidence for pre-existing T cell immunity against SARS-CoV-2, the presence of a pre-existing B cell immunity remains to be elucidated (Grifoni et al., 2020; Le Bert et al., 2020; Mateus et al., 2020; Sette and Crotty, 2021).

Recently published studies that investigated a pre-existing B cell immunity in unexposed individuals provide partially conflicting results. For instance, one study based on the examination of serum samples reported detection of SARS-CoV-2 S protein cross-reactive antibodies and pre-existing humoral immunity in uninfected children, adolescents, or pregnant women (Ng et al., 2020). Cross-reactive humoral immune responses were mainly observed against S protein structures within the S2 subunit that are relatively conserved among endemic HCoVs and SARS-CoV-2. The authors of this study hypothesize that prior immunity against HCoVs may modify COVID-19 disease severity and susceptibility as well as seasonal and geographical transmission patterns. In contrast, other studies only rarely detected cross-reactive serum and B cell responses against SARS-CoV-2 in pre-pandemic samples and argue against a broad and clinically relevant pre-existing B cell immunity (Anderson et al., 2021; Nguyen-Contant et al., 2020; Poston et al., 2020; Shrock et al., 2020; Song et al., 2021). Of note, all studies to date only investigated of plasma/serum and enriched or PBMC secreted IgG fractions of pre-pandemic samples (Anderson et al., 2021; Ng et al., 2020; Nguyen-Contant et al., 2020; Poston et al., 2020; Shrock et al., 2020; Song et al., 2021). They did not involve the analysis of naïve B cell receptor repertoires and putatively SARS-CoV-2-reactive B cells or the characterization of recombinant monoclonal antibodies derived from these cells. Therefore, such plasma-based studies may miss pre-existing potent naïve B cell precursors and low frequency memory B cells which do not contribute sufficient amounts of functionally detectable antibodies to plasma Ig fractions. In our study we cover not only an extensive examination of plasma and IgG fractions but reach out further to investigate the presence of SARS-CoV-2-reactive B cell precursors, memory B cells and the characterization of respective monoclonal antibodies.

We recently isolated SARS-CoV-2-reactive antibodies from convalescent individuals and identified highly similar heavy and/or light chain sequences in naive B cell receptor repertoires from pre-pandemic samples (Kreer et al., 2020a). However, our previous publication did not cover any functional data. Here, we show that some of these chains can replace the original heavy or light chain in SARS-CoV-2-reactive antibodies without altering their functionality. This suggests that the readily detected antibody response in some individuals might use existing antibody heavy or light chains with distinct CDR3 recombination patterns or germline encoded sequence features. In line with this, SARS-CoV-2 neutralizing antibodies were successfully isolated from human naïve antibody gene libraries using phage display (Bertoglio et al., 2021). The most promising antibody candidate STE73-2E9 isolated from these libraries specifically targets the ACE2-RBD interface without cross-reactivity to other coronaviruses and neutralizes authentic SARS-CoV-2 wildtype virus with an IC50 of 0.43 nM. Since phage display relies on random sequence recombination, this study does not provide evidence that SARS-CoV-2-reactive antibodies naturally occur in the naïve B cell repertoire.

Using various immunological and functional assays assessing SARS-CoV-2 binding and neutralization activity as well as cross-reactivity to endemic beta coronaviruses, we found no evidence of significant plasma or IgG reactivity against SARS-CoV-2 in pre-pandemic samples. We comprehensively investigated plasma binding activity of IgG, IgM and IgA immunoglobulin isotypes against diverse beta coronavirus S proteins (SARS-CoV-2 S1 and S1/S2, HCoV-HKU1 and HCoV-OC43 S) by in house and commercially available ELISAs. Moreover, we validated our ELISA results against cell-surface-expressed S protein using flow cytometry and by applying SARS-CoV-2 pseudo- as well as wildtype neutralization assays. Our results are consistent with recently published studies that disagreed on a broad pre-existing B cell background immunity (Anderson et al., 2021; Ng et al., 2020; Nguyen-Contant et al., 2020; Poston et al., 2020; Shrock et al., 2020; Song et al., 2021). However, the findings of our plasma screening may not be directly comparable to the one recent study that indeed reported pre-existing humoral immunity in unexposed children, adolescents, and pregnant women (Ng et al., 2020) since our cohort does not encompass such particular groups of individuals.

To investigate potential pre-existing immunity on a molecular level, we applied antigen-specific single cell sorts to 40 donors and isolated a total of 8,174 putatively SARS-CoV-2-reactive B cells. Consistent with our findings from binding and neutralization screening of plasma samples and polyclonal IgG, frequencies of putatively SARS-CoV-2-reactive B cells in pre-pandemic samples were very low especially when compared to COVID-19 convalescent samples. The choice of an appropriate bait protein for single cell sorting strategies critically determines the isolation of antigen-specific B cells. Immunogenic structures that are not displayed by the chosen bait protein are excluded from isolation. To ensure the comprehensive isolation of antigen-specific B cells, we therefore chose to apply the native, full trimeric SARS-CoV-2 S protein and to adjust the sorting gate. With regard to the low frequencies of SARS-CoV-2-reactive B cells in our study, it can be argued that application of alternative bait proteins e.g. the S2 subunit would have been favorable (Ng et al., 2020; Nguyen-Contant et al., 2020; Shrock et al., 2020; Song et al., 2021). Antibody sequence analyses of isolated putatively SARS-CoV-2-reactive B cells revealed diverse heavy chain V gene usage, normally distributed CDRH3 lengths and V_H_ gene germline identities similar to the naïve BCR receptor repertoire of healthy individuals sampled before the pandemic. In particular, we found no evidence of enrichment of specific V regions which were preferentially used in antibodies of SARS-CoV-2 convalescent individuals such as IGHV3-30, IGHV3-53 or IGHV3-63 (Barnes et al., 2020b, 2020a; Robbiani et al., 2020). Furthermore, we detected no relevant reactivity of 158 recombinantly produced monoclonal antibodies against SARS-CoV-2 and endemic HCoV S proteins.

A pre-existing B cell immunity against pathogens can be a critical determinant of clinical outcome of infections and vaccination strategies. Recent studies provide conflicting results regarding the presence and importance of a pre-existing B cell immunity against SARS-CoV-2 in unexposed individuals. While a pre-existing B cell immunity against SARS-CoV-2 was reported in special study cohorts including children, adolescents or pregnant women, our detailed analysis of the B cell response yielded no comparable evidence in a large population of healthy adults.

## Acknowledgments

We thank all study participants for supporting our research by blood donation; members of the Klein and Becker laboratories for their support and inspiring discussions; Raiees Andrabi, Victor Corman, Jason McLellan for sharing and providing the SARS-CoV-2 and endemic HCoVs S ectodomain plasmids; Daniela Weiland and Nadine Henn for lab management and assistance; Birgit Gathof and Sabine Adam of the Institute of Transfusion Medicine of the University Hospital of Cologne for organizing and providing blood donations of study participants; technical assistants of the serology department at the Institute of Virology of the University Hospital of Cologne for assistance with ELISA testing of plasma samples; as well as Stephan Becker and Verena Kraehling for sharing VeroE6 cells and Susanne Berghöfer for excellent technical assistance with neutralization assays. This work was funded by grants from the German Center of Infection Research (DZIF) to F.K and S.B., the German Research Foundation (DFG) CRC1279 and CRC1310, European Research Council (ERC) ERC-stG639961, and COVIM: NaFoUniMed-Covid19 to F. Klein.

## Author Contributions

Conceptualization – F.K., M.S.E., L.G., C.K. and M.Z.; Methodology – M.S.E., L.G., S. Dähling, N.P., S. Dettmer, M. Korenkov, K.V., V.D.C., S.H, Michael Klüver, C.K.; Formal analysis – M.S.E., L.G., C.K., V.D.C, S.H., V.K., S. Dähling; Investigation – M.S.E., L.G., C.K., V.D.C., S.H., V.K., S. Dähling, S. Dettmer, M. Korenkov, and N.P.; Resources – B.G.; Writing original draft - M.S.E., L.G., C.K. and F.K.; Writing reviewing and editing – all authors; Supervision – F.K., C.K.; Funding acquisition – F.K..

These authors contributed equally to the work: M.S.E. and L.G.

## STAR Methods

### RESOURCE AVAILABILITY

#### Lead contact

Further information and requests for resources and reagents should be directed to and will be fulfilled by the Lead Contact, Florian Klein.

#### Materials availability

There are restrictions to the availability of SARS-CoV-2-reactive antibodies due to limited production capacities and ongoing consumption. Reasonable amounts of antibodies will be made available by the Lead Contact upon request under a Material Transfer Agreement (MTA) for non-commercial usage. Nucleotide sequences and expression plasmids will be shared upon request.

#### Data and code availability

Nucleotide sequences of isolated monoclonal antibodies will be deposited at GenBank after review or acceptance of the manuscript. Further heavy and light chain sequences as well as NGS data of healthy individuals will be shared by the Lead Contact upon request.

### EXPERIMENTAL MODEL AND SUBJECT DETAILS

#### Study participants and collection of clinical samples

Study participants were recruited at the University Hospital Cologne. Buffy coats from study participants were collected according to a study protocol approved by the Institutional Review Board of the University of Cologne (study protocol 16-054). Buffy coats were provided as residual products in the context of regular blood donations from the Institute for Transfusion Medicine at the University Hospital of Cologne. All study participants provided informed consent. The study cohort comprised 150 individuals with 50.7% female and 49.3% male adults. All study participants were ≥ 18 years old. Blood samples were collected between August and November 2019.

#### Cell lines

HEK293T cells were maintained in Dulbecco’s Modified Eagle Medium (DMEM, Thermo Fisher) supplemented with 10 % fetal bovine serum (FBS, Sigma-Aldrich), 1 x antibiotic-antimycotic (Thermo Fisher), 1 mM sodium pyruvate (Gibco) and 2 mM L-glutamine (Gibco) at 37 °C and 5 % CO_2_. HEK293-6E cells were maintained in FreeStyle 293 Expression Medium (Life Technologies) supplemented with 0.2 % penicillin/streptomycin under constant shaking at 90 – 120 rpm, 37 °C and 6 % CO_2_. VeroE6 and HEK293T-ACE2 cells were maintained in Dulbecco’s Modified Eagle Medium (Gibco) supplemented with 10 % fetal bovine serum (FBS, Sigma-Aldrich), 1 x penicillin-streptomycin (Gibco), 1 mM sodium pyruvate (Gibco) and 2 mM L-glutamine (Gibco) at 37 °C and 5 % CO_2_. The sex of HEK293T, HEK293-6E and VeroE6 cell lines is female. Cell lines were not specifically authenticated.

### METHOD DETAILS

#### Isolation of PBMCs, plasma and total IgG

Peripheral blood mononuclear cells (PBMCs) were isolated from buffy coats and whole blood by density gradient separation using Histopaque separation medium (Sigma-Aldrich) and Leucosep cell tubes (Greiner Bio-one) according to the manufacturer’s instructions. Isolated PBMCs were cryopreserved at −150 °C in 90 % FBS supplemented with 10 % DMSO until further use. Plasma was collected and stored at −80 °C. Plasma samples were heat-inactivated at 56 °C for 40 min prior to further use. For IgG isolation, 1 ml of heat-inactivated plasma was incubated with Protein G Sepharose (GE Life Sciences) overnight at 4 °C (GE Life Sciences) under constant rotation. Protein G beads were transferred to chromatography columns and washed twice with sterile PBS. IgGs were eluted from columns using 0.1 M glycine (pH = 3.0) and buffered in 0.1 M Tris (pH = 8.0). Buffer exchange to PBS and concentration of IgGs was performed by centrifugation using 30 kDa Amicon spin membranes (Millipore). The final concentration of purified IgGs was determined by UV/Vis spectroscopy using a Nanodrop (A280). Subsequently, purified IgGs were stored at 4 °C.

#### Expression and purification of viral surface proteins

Constructs encoding the stabilized spike protein ectodomain of the endemic HCoV-HKU1 (amino acids 1 – 1295, GenBank ID.: ABD75497) and HCoV-OC43 (amino acids 1 – 1300, GenBank ID.: AAX84792) were kindly provided by Raiees Andrabi (California, USA) and Victor Corman (Berlin, Germany) and described previously (St-Jean et al., 2004; Woo et al., 2005; Corman et al., 2012; Kreye et al., 2020; Song et al., 2021). Constructs encoding the prefusion stabilized SARS-CoV-2 S ectodomain (amino acids 1 – 1207; GenBank ID.: MN908947) were kindly provided by Jason McLellan (Texas, USA) and Florian Krammer (New York, USA) and previously described (Stadlbauer et al., 2020; Wrapp et al., 2020). All three constructs contain the following mutations: two proline substitutions for prefusion state stabilization (SARS-CoV-2: residues 986 and 987; OC43: residues 1078 and 1079; HKU1: residues 1071 and 1072) and the furin cleavage sites were mutated (SARS-CoV-2: ‘‘GGGG’’ substitution at residues 682–685; OC43: ‘‘GSAS’’ substitution at residues 762-766; HKU1‘‘GSAS’’ substitution at residues 756-760). Ebola surface glycoprotein (EBOV Makona, GenBank ID.: KJ660347; amino acids 1 – 651) was used as a negative control for HCoV S protein ELISAs. The Ebola surface glycoprotein was stabilized with GCN4 trimerization domain and expressed without the transmembrane domain as previously described (Ehrhardt et al., 2019). The different coronavirus ectodomains were amplified from the synthetic gene plasmids by PCR and subsequently cloned into a modified sleeping beauty transposon expression vector containing a C-terminal T4 fibritin trimerization motif (foldon) followed by a Twin-Strep-Tag purification tag. For the recombinant protein productions, stable HEK293 EBNA cell lines were generated employing the sleeping beauty transposon system (Kowarz et al., 2015). Briefly, expression constructs were transfected into the HEK293 EBNA cells using FuGENE HD transfection reagent (Promega). After selection with puromycin, cells were induced with doxycycline. Cell supernatants were filtered and the recombinant proteins purified via Strep-Tactin XT (IBA Lifescience) resin. Proteins were then eluted by biotin-containing TBS-buffer (IBA Lifescience), and dialyzed against TBS-buffer.

#### Isolation of single SARS-CoV-2-reactive B cells

CD19^+^ B cells were enriched from PBMCs by immunomagnetic cell separation using CD19 microbeads (Miltenyi Biotec) according to the manufacturer’s protocol. Enriched B cells were labeled for 20 min on ice with 4’,6-Diamidin-2-phenlindol (DAPI; Thermo Fisher Scientific), anti-human CD20-Alexa Fluor 700 (BD), anti-human IgG-APC (BD), anti-human CD27-PE (BD) and DyLight488-labeled SARS-CoV-2 S protein (10 μg/ml). DAPI^−^, CD20^+^, SARS-CoV S protein^+^, IgG^+^/IgG^−^ single cells were sorted into 96 well plates using a FACSAria Fusion (Becton Dickinson). Each well of the 96 well plate was prefilled with 4 μl of sorting buffer consisting of 0.5 x PBS, 0.5 U/μl RNAsin (Promega), 0.5 U/μl RNaseOut (Thermo Fisher Scientific), and 10 mM DTT (Thermo Fisher Scientific). Plates were cryopreserved at −80 °C immediately after sorting.

#### B cell receptor amplification and sequence analysis

Generation of cDNA and amplification of antibody heavy and light chain genes from sorted single cells was performed as previously described (Gieselmann et al., 2021; Kreer et al., 2020a; Schommers et al., 2020). For reverse transcription, sorted cells were incubated with 7 μl of a random-hexamer-primer master mix (RHP mix) consisting of 0.75 μl Random Hexamer Primer (Thermo Fisher Scientific), 0.5 μl NP-40 (Thermo Fisher Scientific), 0.15 μl RNaseOut (Thermo Fisher Scientific), and 5.6 μl of nuclease-free H_2_O at 65 °C for 1 min. Subsequently, samples were incubated with a reverse-transcription master mix (RT mix) consisting of 3 μl of 5 x Superscript IV RT buffer (Thermofisher Scientific), 0.5 μl dNTPs (Thermo Fisher Scientific), 1 μl DTT (Thermo Fisher Scientific), 0.1 μl of RNasin (Promega), 0.1 μl of RNaseOut (Thermo Fisher Scientific), 2.05 μl of nuclease-free H_2_O and 0.25 μl of Superscript IV (Thermofisher Scientific) per well and incubated at 42 °C for 10 min, 25 °C for 10 min, 50°C for 10 min and 94 °C for 5 min. Heavy and light chains were amplified from cDNA by semi-nested, single cell PCRs using optimized V gene-specific primer mixes(Kreer et al., 2020b) and Platinum Taq DNA Polymerase or Platinum Taq Green Hot Start Polymerase (Thermo Fisher Scientific) as previously described (Kreer et al., 2020a; Schommers et al., 2020; Kreer et al., 2020b; Gieselmann et al., 2021). PCR products were analyzed by agarose gel electrophoresis for correct product size and subsequently send for Sanger sequencing. Only chromatograms with a mean Phred score of 28 and sequences with a minimal length of 240 nucleotides were selected for downstream sequence analyses. Filtered sequences were annotated with IgBlast (Ye et al., 2013) according to the IMGT system (Lefranc et al., 2009) and only the variable region from FWR1 to the end of the J gene was extracted. Base calls within the variable region with a Phred score below 16 were masked and sequences with more than 15 masked nucleotides, stop codons, or frameshifts were excluded from further analyses. Sequence analyses to inform on sequence clonality were performed separately for each study participant. All productive heavy chain sequences were grouped by identical V_H_/J_H_ gene pairs and the pairwise Levenshtein distance for their CDRH3s was determined. Starting from a random sequence, clone groups were assigned to sequences with a minimal CDRH3 amino acid identity of at least 75 % (with respect to the shortest CDRH3). 100 rounds of input sequence randomization and clonal assignment were performed and the result with the lowest number of remaining unassigned (non-clonal) sequences was selected for downstream analyses. All clones were cross-validated by the investigators taking shared mutations and light chain information into account.

#### Sequence selection for antibody production

Antibody selection for cloning was done by two approaches. First, a similarity search was performed against 868 and 52 SARS-CoV-2-binding antibodies from the CoVAbDab (Raybould et al., 2020) (retrieved on 21.08.20) and Kreer et al 2020 (Kreer et al., 2020a), respectively. B cell receptor sequences with identical VH/JH combination, a CDRH3 length difference ≤ 2 AA, and a CDRH3 Levenshtein distance ≤ 3 AA in comparison to at least one SARS-CoV-2-binding antibody were selected as similar (18 in total). Second, a random selection was performed to yield at least 6 random antibody sequences per individual including at least 3 different clones and at least 3 non-clonal sequences. Random non-clonal sequences were used to fill up the selection if less than 3 clones were available.

#### Cloning and production of monoclonal antibodies

Heavy and light chain variable regions of selected antibodies were cloned into expression vectors by sequence and ligation independent cloning (SLIC) (von Boehmer et al., 2016) as previously described (Tiller et al., 2008; Schommers et al., 2020; Kreer et al., 2020a; Gieselmann et al., 2021). 1^st^ PCR product was amplified using Q5 Hot Start High Fidelity DNA Polymerase (New England Biolabs) and specific forward and reverse primers including adaptor sequences which are homologous to the restriction sites of the antibody expression vector (IgG1, IgL, IgK (Tiller et al., 2008)). Forward primers were designed according to 2^nd^ PCR primers (Kreer et al., 2020b) and encode for the complete native leader sequence of all heavy and light chain V genes, whereas reverse primers bind to the conserved sequence motifs at the 5’ end of heavy and light chain immunoglobulin constant regions. PCR was run at 98 °C for 30 s; 35 cycles of 98 °C for 10 s, 72 °C for 45 s; and 72 °C for 2 min. Subsequently, PCR products were purified with 96-well format silica membranes, cloned by SLIC (von Boehmer et al., 2016) into linearized expression vectors with T4 DNA polymerase (NEB) and transformed into chemically competent *Escherichia coli* (DH5α). Heavy chain variable regions of IgG^−^ B cells were cloned into IgG1 expression vectors. Correct insertion of the V region sequence into the expression vector was examined by colony PCR and Sanger sequencing. Positive colonies were propagated in midi cultures and plasmids purified.

For production of recombinant monoclonal antibodies, HEK293-6E suspension cells were co-transfected chemically with human heavy chain and corresponding light chain antibody expression vectors using 25 kDa branched polyethylenimine (PEI; Sigma-Aldrich). Transfected cells were propagated in FreeStyle 293 Expression Medium (Thermo Fisher Scientific) supplemented with 0.2 % penicillin/streptomycin (Thermo Fisher Scientific) at 37 °C and 6 % CO_2_ under constant shaking at 90 – 120 rpm for seven days. Cell supernatants were harvested by centrifugation, filtered through PES filters and incubated overnight at 4 °C under constant rotation with Protein-G-coupled Sepharose beads (GE Life Sciences). The suspension was transferred to chromatography columns, washed twice with sterile PBS and antibodies were eluted with 0.1 M glycine (pH = 3) and buffered using 1 M Tris (pH = 8). Buffer exchange to PBS and concentration of antibodies was performed by centrifugation using 30kDa Amicon spin membranes (Millipore). Antibodies were filtered through Ultrafree-MC 0.22 μm membranes (Millipore) and stored at 4 °C. The final concentration of purified antibodies was determined by UV/Vis spectroscopy using a Nanodrop (A280). Subsequently, purified antibodies were stored at 4 °C.

#### Cell-surface-expressed S protein immunoassay

HEK293T cells were transfected with plasmids encoding the full-length SARS-CoV-2 S protein (GenBank ID.: MN908947) using TurboFect™ transfection reagent (Thermo Fisher Scientific). After incubation at 37 °C and 5 % CO_2_ for 48 h, adherent cells were detached with PBS supplemented with 1 mM EDTA (pH = 7.4) and resuspended in FACS buffer (1x PBS supplemented with 2 % FCS and 2 mM EDTA). 3 x 10^4^ cells were distributed in 50 μl of FACS buffer to each well of a V-bottom-shaped 96 well plate. Starting with an initial dilution of 1:25, plasma samples were prepared in a 4-fold serial dilution for a total of 6 dilutions. Cells were incubated with 50 μl/well of diluted plasma samples and incubated for 30 min on ice. After incubation, cells were washed once with 100 μl/well of FACS buffer and stained in 50 μl/well of 1:160 dilution of BV421 IgG and 1:100 dilution of APC IgM on ice for 30 min. After staining, cells were washed once with 100 μl/well of FACS buffer and analyzed on a FACS BD Aria Fusion. Evaluations were performed using FlowJo10 software. Geometric Mean values of all cells/single cells/mCherry positive (transfected cells) in APC-A channel (for IgM) and BV421-A channel (for IgG) were determined and graphically displayed using GraphPad Prism.

#### SARS-CoV-2 and HCoV S protein immunoassays

ELISA plates (Greiner Bio-One 655092) were coated with 2 μg/ml protein (spike ectodomains of SARS-CoV-2, HKU1, OC43) for IgG measurement or 5 μg/ml (spike ectodomain of SARS-CoV-2) for IgM and IgA measurements in PBS at 4 °C overnight. Plates were blocked with blocking buffer (BB) consisting of PBS supplemented with 5 % nonfat dried milk powder (Carl Roth T145.2) for 60 min at RT. Monoclonal antibodies were tested with a starting concentration of 10 μg/ml in PBS, polyclonal IgGs with 500 μg/ml in BB and plasma samples with a starting dilution of 1:20 for IgG detection and 1:10 for IgM and IgA detection in BB. Monoclonal antibodies were serially diluted 1:5. Polyclonal IgGs and plasma were diluted in 1:4 serial dilutions. After incubation with samples for 90 min at RT, plates were incubated with anti-human IgG-HRP (Southern Biotech 2040-05) diluted 1:2500 in BB, anti-human IgM-HRP (Thermo Fisher Scientific A18835) diluted 1:2000 in BB or anti-human IgA-HRP (Thermo Fisher Scientific A18781) 1:2000 in BB. Plates were developed using ABTS solution (Thermo Fisher Scientific 002024) and absorbance was measured at 415 nm and 695 nm by a Microplate Reader (Tecan).

Anti-SARS-CoV-2 S1/S2 IgG and IgM antibody titers of plasma samples were also assessed using the automated DiaSorin’s LIAISON® SARS-CoV-2 S1/S2 protein ELISA kit according to the manufacturer’s instructions. IgG and IgM result values were interpreted with the following cut-off values: negative < 12.0 AU/ml, equivocal ≥ 12.0 - < 15.0 AU/ml, and positive ≥ 15.0 AU/ml. Anti-SARS-CoV-2 S1 IgA antibody titers of plasma samples were also measured using the Euroimmun anti-SARS-CoV-2 ELISA on the automated system Euroimmun Analyzer I and S/CO values interpreted with following cut-off values: negative S/CO < 0.8, equivocal S/CO ≥ 0.8 - < 1.1, positive S/CO ≥ 1.1.

#### SARS-CoV-2 pseudovirus neutralization assays

SARS-CoV-2 pseudovirus expressing the Wu01 spike (EPI_ISL_40671) was generated by co-transfection of individual plasmids encoding HIV Tat, HIV Gag/Pol, HIV Rev, luciferase followed by an IRES and ZsGreen, and the SARS-CoV-2 spike protein into HEK 293T cells using the FuGENE 6 Transfection Reagent (Promega). Cell culture supernatants containing pseudovirus particles were harvested and stored at −80°C till use. The pseudovirus was titrated by infecting HEK293T cells expressing human ACE2 (Crawford et al., 2020). Following a 48-hour incubation at 37°C and 5% CO_2_, luciferase activity was determined by addition of luciferin/lysis buffer (10 mM MgCl2, 0.3 mM ATP, 0.5 mM Coenzyme A, 17 mM IGEPAL (all Sigma-Aldrich), and 1 mM D-Luciferin (GoldBio) in Tris-HCL) using a microplate reader (Berthold). For neutralization assays, a virus dilution with a relative luminescence unit (RLU) of approximately 1000-fold in infected cells versus non-infected cells was selected.

For testing neutralization at a single dilution, polyclonal IgG samples at a concentration of 1000 μg/ml, plasma samples at a dilution of 1:10, or mAbs at a concentration of 50 μg/ml, were co-incubated with pseudovirus supernatants for 1 h at 37°C, following which 293T-ACE-2 cells were added. After a 48 h incubation at 37 °C and 5 % CO_2_, luciferase activity was determined using the luciferin/lysis buffer. After subtracting background RLUs of non-infected cells, % of neutralization was calculated and the mean value was used for reporting. Each sample was tested in duplicates.

To determine IC_50_ values for mAbs a dilution series of the antibody was performed starting with 50 μg/ml. IC_50_ values were calculated as the antibody concentration causing a 50 % reduction in signal compared the virus-only controls using a dose-response curve in GraphPad Prism.

#### Authentic virus neutralization assays

Authentic virus neutralization was tested using a virus previously grown out from an oro-/naso-pharyngeal swab using VeroE6 cells (Vanshylla et al., 2021). For testing neutralization, plasma samples at a single dilution of 1:10 were co-incubated with the 200 TCID_50_ virus for 1 h at 37°C, following which VeroE6 cells were added. After 4 days, cytopathic effects (CPE) were analysed under a bright-field microscope and neutralization was determined as the absence of CPE. Cells without any virus served as reference for lack of CPE and cells with virus only served as reference for positive CPE.

### QUANTIFICATION AND STATISTICAL ANALYSIS

Flow cytometry analyses and quantification were performed using FlowJo10 software. Statistical tests and analyses were done with GraphPad Prism (v7 and v8), Python (v3.6.8), R (v4.0.0) and Mircosoft Excel for Mac (v14.7.3 and 16.4.8). CDRH3 lengths, V gene usage and germline identity distributions for clonal sequences (**Figure 3C**) were assessed for all input sequences without further collapsing. A two-tailed unpaired t test (Prism, GraphPad) was performed to test for the frequency differences of SARS-CoV-2-reactive IgG+ and IgG- B cells between pre-pandemic and convalescent blood samples (**Figure 3A**).

### KEY RESOURCES TABLE

### Supplementary Figure Legends

**Figure S1: Binding of plasma samples and polyclonal IgG, related to Figure 2**

Binding of plasma IgG, IgM and IgA to SARS-CoV-2 S protein determined by serial dilution **(A)** or automated ELISAs **(B)** (DiaSorin’s LIAISON® and Euroimmun Analyzer). **(C)** Binding of purified, polyclonal IgGs (pIgG) to SARS-CoV-2, HKU1 and OC43 S proteins determined by serial dilution ELISA. ELISAs were performed in duplicate experiments. Circles depict means and error bars indicate standard deviation. IgG titers were interpreted as negative < 12.0 AU/ml, equivocal ≥ 12.0 - < 15.0 AU/ml, and positive ≥ 15.0 AU/ml. IgM titers were interpreted as negative < 1.1 IgM index and positive > 1.1. IgA values interpreted with following cut-off values: negative < 0.8 IgA ratio, equivocal ≥ 0.8 - < 1.1 IgA ratio, positive ≥ 1.1 IgA ratio.

**Figure S2: Neutralization activity of plasma and polyclonal IgGs against SARS-CoV-2, related to Figure 2**

Neutralization activity of plasma samples **(A)** and polyclonal IgGs **(B)** against SARS-CoV-2 wildtype (WT) and/or pseudovirus (PSV). Samples were tested in duplicate experiments (wildtype) or in duplicates in a single experiment (pseudovirus). The SARS-CoV-2 neutralizing antibody C6 was used as a positive control. Bars and circles of graph plots show means and error bars indicate standard deviation.

**Figure S3: Gating strategy and single cell sorts of SARS-CoV-2-reactive B cell subsets, related to Figure 3**

**(A)** FACS plots illustrating the gating strategy for single cell sorts of SARS-CoV-2-reactive, IgG^+^ and IgG^−^ B cells. **(B)** Individual FACS plots depicting sorting gates and frequencies of SARS-CoV-2-reactive, IgG^+^ and IgG^−^ B cells from 40 donors.

**Figure S4: Binding of monoclonal antibodies isolated from pre-pandemic blood samples, related to Figure 4**

**(A)** Binding of monoclonal antibodies to SARS-CoV-2, HKU-1 and OC43 S proteins determined by serial dilution ELISA. ELISAs were performed in duplicate experiments. Circles depict means and error bars indicate standard deviation.

### Supplementary Tables

**Table S1: Demographical characteristics of blood donors, related to Figure 2**

**Table S2: Sequence information of pre-pandemic and original heavy and light chains, related to Figure 1**

