## Supplementary information for "No substantial pre-existing B cell immunity against SARS-CoV-2 in healthy adults"

### Supplementary Figures


| | |
|---|---|
| Figure S1 | Binding of plasma samples and polyclonal IgG |
| Figure S2 | Neutralization activity of plasma and polyclonal IgGs against SARS-CoV-2 |
| Figure S3 | Gating strategy and single cell sorts of SARS-CoV-2-reactive B cell subsets |
| Figure S4 | Binding of monoclonal antibodies isolated from pre-pandemic blood samples |


### Supplementary Tables


| | |
|---|---|
| Table S1 | Demographical characteristics of blood donors |
| Table S2 | Sequence information of pre-pandemic and original heavy and light chains |

**Figure S1**

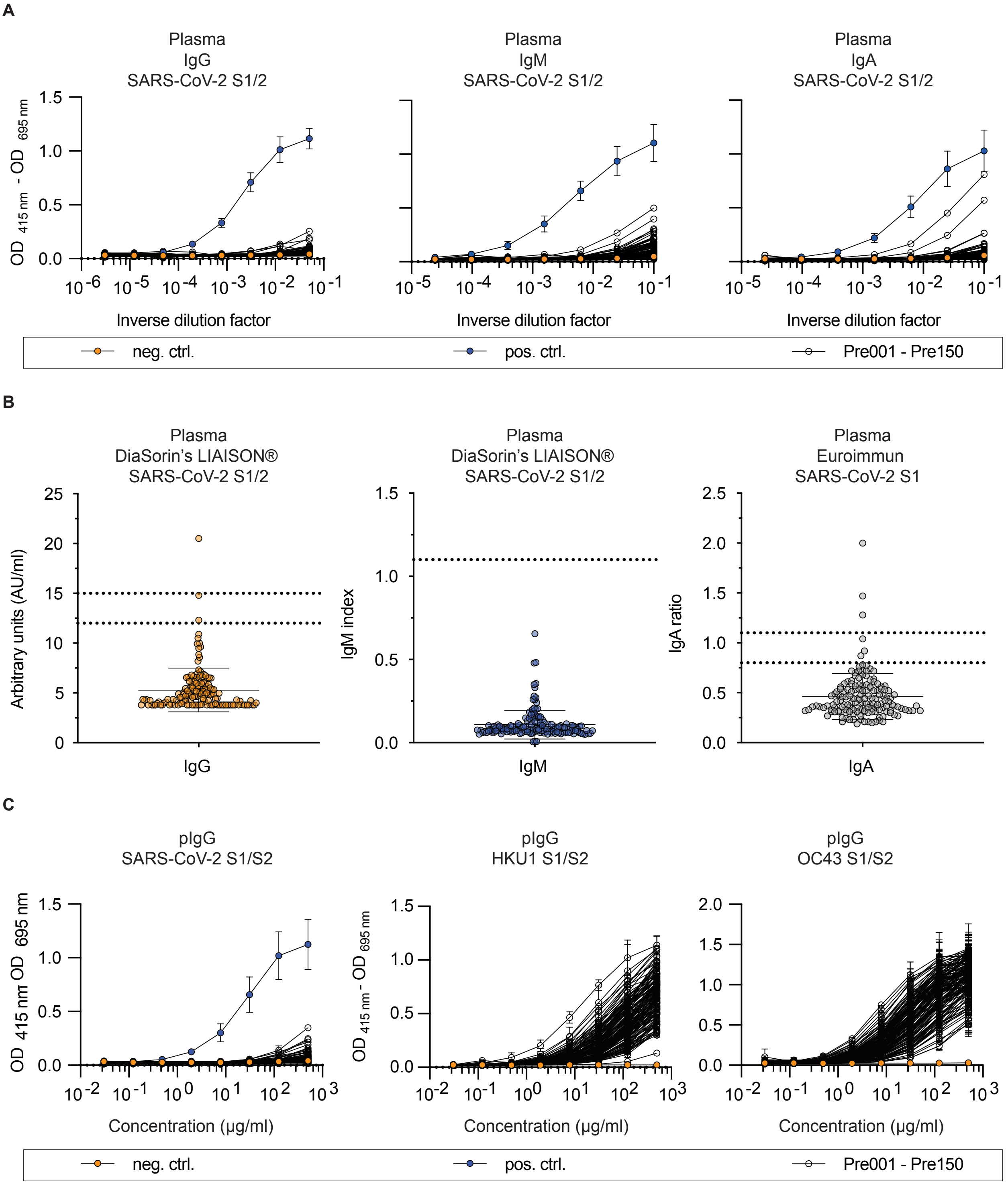


**Figure S1: Binding of plasma samples and polyclonal IgG, related to Figure 2**
Binding of plasma IgG, IgM and IgA to SARS-CoV-2 S protein determined by serial dilution (A) or automated immunoassay (B) (DiaSorin's LIAISON® and Euroimmun Analyzer). (C) Binding of purified, polyclonal IgGs (pIgG) to SARS-CoV-2, HKU1 and OC43 S proteins determined by serial dilution ELISA. ELISAs were performed in duplicate experiments. Circles depict means and error bars indicate standard deviation. IgG titers were interpreted as negative < 12.0 AU/ml, equivocal > 12.0 - < 15.0 AU/ml, and positive > 15.0 AU/ml. IgM titers were interpreted as negative < 1.1 IgM index and positive > 1.1. IgA values interpreted with following cut-off values: negative < 0.8 IgA ratio, equivocal > 0.8 - < 1.1 IgA ratio, positive > 1.1 IgA ratio.

**Figure S2**
**A**

**Plasma.** SARS-CoV-2 wildtype neutralization assay

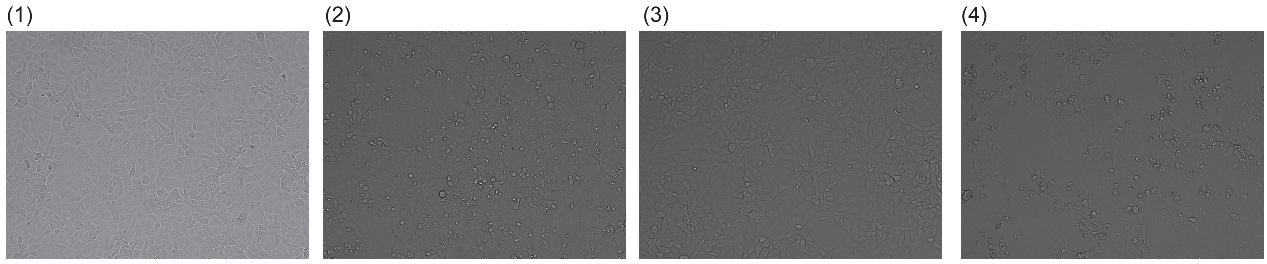


(1) uninfected VeroE6 cells after 96 hours, no sign of cytopathic effect (CPE).
(2) WT SARS-CoV-2 infected VeroE6 cells after 96 hours, with visible CPE.
(3) WT SARS-CoV-2 infected VeroE6 cells after 96 hours in the presence of SARS-CoV-2 neutralizing mAb C6, no sign of CPE.
(4) WT SARS-CoV-2 infected VeroE6 cells after 96 hours in the presence of 1:10 diluted plasma from a prepandemic sample, with visible CPE. Representative result for all 150 tested prepandemic plasma samples with clear sign of CPE among all tested samples.

**Plasma.** SARS-CoV-2 PSV neutralization assay

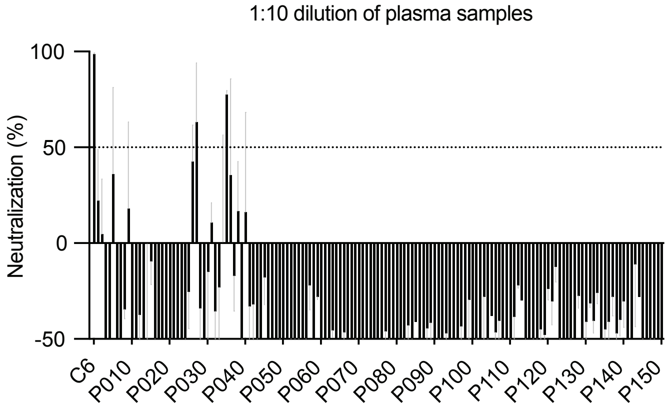


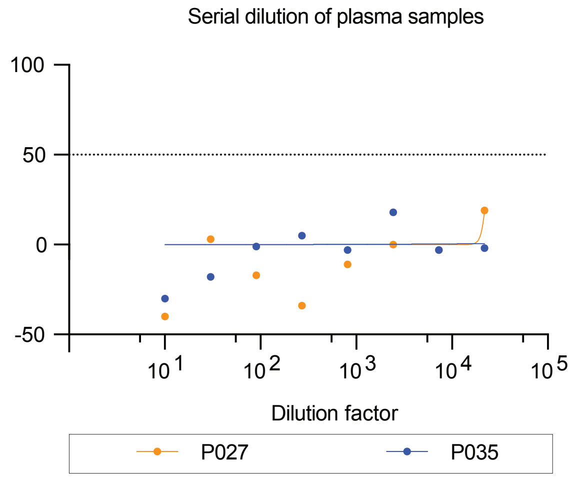


**B**

**Polyclonal IgGs.** SARS-CoV-2 PSV neutralization assay

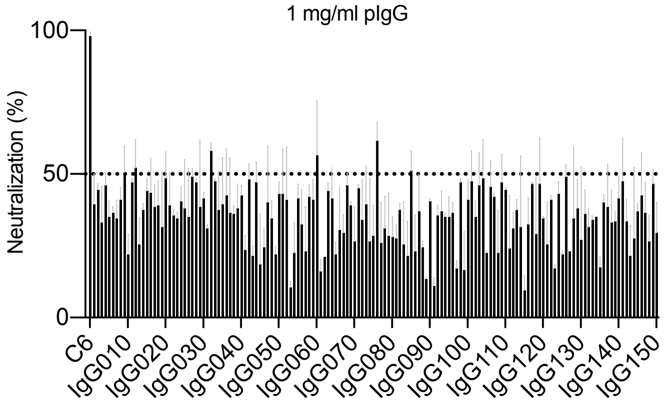


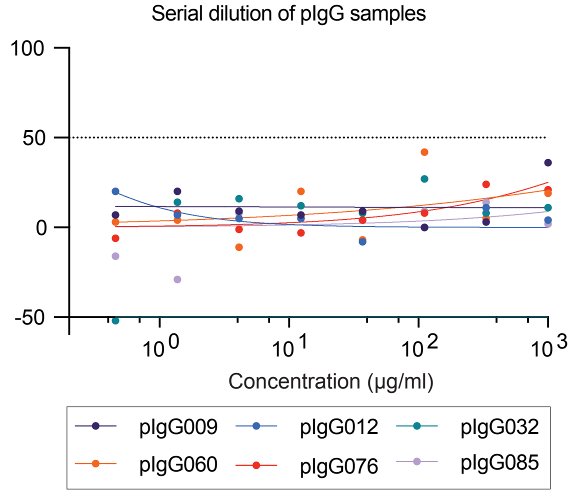


**Figure S2: Neutralization activity of plasma and polyclonal IgGs against SARS-CoV-2, related to Figure 2**
Neutralization activity of plasma samples (A) and polyclonal IgGs (B) against SARS-CoV-2 wildtype (WT) and/or pseudovirus (PSV). Samples were tested in duplicate experiments (wildtype) or in duplicates in a single experiment (pseudovirus). The SARS-CoV-2 neutralizing antibody C6 was used as a positive control. Bars and circles of graph plots show means and error bars (light gray) indicate standard deviation. Samples tested in single dilution exceeding neutralization activity > 50 % were repeatedly tested in serial dilutions.

**Figure S3**

A

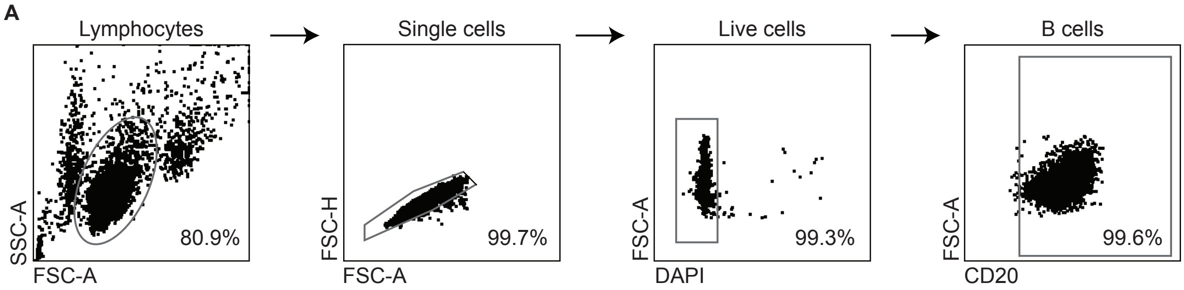
Lymphocytes
Single cells
Live cells
B cells
SSC-A
FSC-A
80.9%
FSC-H
FSC-A
99.7%
FSC-A
DAPI
99.3%
FSC-A
CD20
99.6%

B

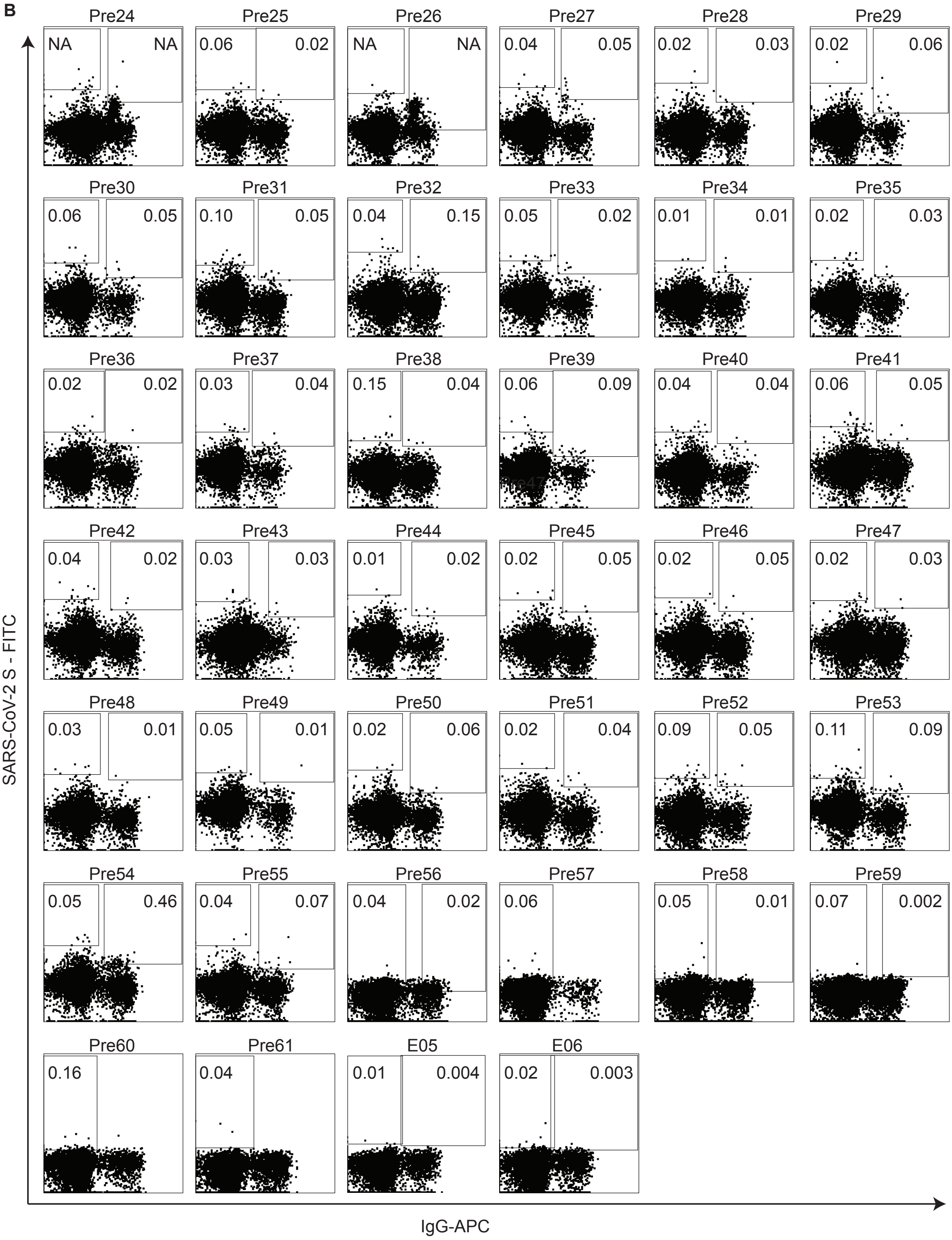
Pre24 NA NA
Pre25 0.06 0.02
Pre26 NA NA
Pre27 0.04 0.05
Pre28 0.02 0.03
Pre29 0.02 0.06
Pre30 0.06 0.05
Pre31 0.10 0.05
Pre32 0.04 0.15
Pre33 0.05 0.02
Pre34 0.01 0.01
Pre35 0.02 0.03
Pre36 0.02 0.02
Pre37 0.03 0.04
Pre38 0.15 0.04
Pre39 0.06 0.09
Pre40 0.04 0.04
Pre41 0.06 0.05
Pre42 0.04 0.02
Pre43 0.03 0.03
Pre44 0.01 0.02
Pre45 0.02 0.05
Pre46 0.02 0.05
Pre47 0.02 0.03
Pre48 0.03 0.01
Pre49 0.05 0.01
Pre50 0.02 0.06
Pre51 0.02 0.04
Pre52 0.09 0.05
Pre53 0.11 0.09
Pre54 0.05 0.46
Pre55 0.04 0.07
Pre56 0.04 0.02
Pre57 0.06
Pre58 0.05 0.01
Pre59 0.07 0.002
Pre60 0.16
Pre61 0.04
E05 0.01 0.004
E06 0.02 0.003
SARS-CoV-2 S - FITC
IgG-APC

**Figure S3: Gating strategy and single cell sorts of SARS-CoV-2-reactive B cell subsets, related to Figure 3**
(A) FACS plots illustrating the gating strategy for single cell sorts of SARS-CoV-2-reactive, IgG+ and IgG- B cells.
(B) Individual FACS plots depicting sorting gates and frequencies of SARS-CoV-2-reactive, IgG+ and IgG- B cells from 40 donors.

**Figure S4**

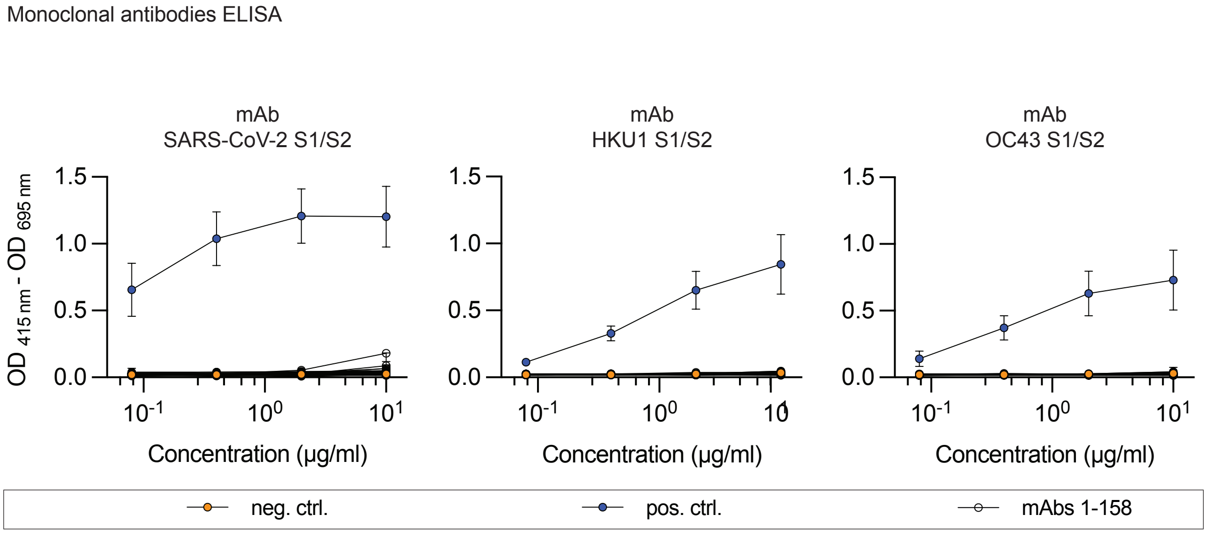


**Figure S4: Binding of monoclonal antibodies isolated from pre-pandemic blood samples, related to Figure 4**
Binding of monoclonal antibodies to SARS-CoV-2, HKU-1 and OC43 S proteins determined by serial dilution ELISA. ELISAs were performed in duplicate experiments. Circles depict means and error bars indicate standard deviation.

**Table S1: Demographical characteristics of blood donors, related to Figure 2**

| Sample | Date of blood draw | Age | Gender | Sample | Date of blood draw | Age | Gender | Sample | Date of blood draw | Age | Gender |
|---|---|---|---|---|---|---|---|---|---|---|---|
| Pre001 | 08.08.2019 | 26 | w | Pre052 | 16.10.2019 | 28 | w | Pre103 | 24.10.2019 | 34 | m |
| Pre002 | 08.08.2019 | 28 | w | Pre053 | 16.10.2019 | 38 | m | Pre104 | 25.10.2019 | 32 | w |
| Pre003 | 08.08.2019 | 26 | w | Pre054 | 16.10.2019 | 41 | w | Pre105 | 25.10.2019 | 23 | w |
| Pre004 | 08.08.2019 | 26 | w | Pre055 | 16.10.2019 | 22 | w | Pre106 | 25.10.2019 | 23 | w |
| Pre005 | 08.08.2019 | 36 | w | Pre056 | 16.10.2019 | 29 | m | Pre107 | 25.10.2019 | 23 | w |
| Pre006 | 08.08.2019 | 45 | m | Pre057 | 16.10.2019 | 23 | w | Pre108 | 25.10.2019 | 26 | m |
| Pre007 | 08.08.2019 | 61 | m | Pre058 | 16.10.2019 | 23 | w | Pre109 | 25.10.2019 | 26 | m |
| Pre008 | 08.08.2019 | 23 | w | Pre059 | 16.10.2019 | 21 | w | Pre110 | 25.10.2019 | 33 | w |
| Pre009 | 08.08.2019 | 37 | m | Pre060 | 17.10.2019 | 23 | w | Pre111 | 25.10.2019 | 32 | m |
| Pre010 | 08.08.2019 | 56 | m | Pre061 | 17.10.2019 | 26 | m | Pre112 | 25.10.2019 | 42 | m |
| Pre011 | 08.08.2019 | 32 | m | Pre062 | 17.10.2019 | 22 | w | Pre113 | 07.11.2019 | 36 | m |
| Pre012 | 07.10.2019 | 23 | m | Pre063 | 17.10.2019 | 22 | w | Pre114 | 07.11.2019 | 36 | m |
| Pre013 | 07.10.2019 | 30 | m | Pre064 | 17.10.2019 | 22 | w | Pre115 | 07.11.2019 | 24 | w |
| Pre014 | 07.10.2019 | 23 | m | Pre065 | 17.10.2019 | 42 | w | Pre116 | 07.11.2019 | 22 | w |
| Pre015 | 07.10.2019 | 31 | w | Pre066 | 17.10.2019 | 28 | m | Pre117 | 07.11.2019 | 23 | w |
| Pre016 | 09.10.2019 | 20 | w | Pre067 | 17.10.2019 | 28 | m | Pre118 | 07.11.2019 | 24 | m |
| Pre017 | 09.10.2019 | 30 | m | Pre068 | 17.10.2019 | 21 | w | Pre119 | 07.11.2019 | 22 | m |
| Pre018 | 09.10.2019 | 19 | m | Pre069 | 17.10.2019 | 21 | m | Pre120 | 07.11.2019 | 43 | m |
| Pre019 | 09.10.2019 | 28 | w | Pre070 | 17.10.2019 | 23 | w | Pre121 | 07.11.2019 | 24 | m |
| Pre020 | 09.10.2019 | 27 | w | Pre071 | 17.10.2019 | 24 | m | Pre122 | 07.11.2019 | 23 | w |
| Pre021 | 09.10.2019 | 39 | m | Pre072 | 17.10.2019 | 29 | m | Pre123 | 07.11.2019 | 31 | w |
| Pre022 | 09.10.2019 | 22 | w | Pre073 | 18.10.2019 | 19 | w | Pre124 | 07.11.2019 | 33 | m |
| Pre023 | 09.10.2019 | 37 | m | Pre074 | 18.10.2019 | 25 | w | Pre125 | 08.11.2019 | 51 | m |
| Pre024 | 10.10.2019 | 36 | m | Pre075 | 18.10.2019 | 26 | w | Pre126 | 08.11.2019 | 25 | w |
| Pre025 | 10.10.2019 | 29 | m | Pre076 | 18.10.2019 | 27 | w | Pre127 | 08.11.2019 | 66 | w |
| Pre026 | 10.10.2019 | 27 | m | Pre077 | 18.10.2019 | 66 | m | Pre128 | 08.11.2019 | 26 | w |
| Pre027 | 10.10.2019 | 29 | m | Pre078 | 18.10.2019 | 29 | w | Pre129 | 08.11.2019 | 25 | w |
| Pre028 | 10.10.2019 | 22 | m | Pre079 | 22.10.2019 | 20 | w | Pre130 | 12.11.2019 | 19 | w |
| Pre029 | 10.10.2019 | 23 | w | Pre080 | 22.10.2019 | 22 | m | Pre131 | 12.11.2019 | 30 | m |
| Pre030 | 10.10.2019 | 25 | w | Pre081 | 22.10.2019 | 25 | w | Pre132 | 12.11.2019 | 30 | m |
| Pre031 | 10.10.2019 | 20 | w | Pre082 | 22.10.2019 | 57 | w | Pre133 | 12.11.2019 | 47 | w |
| Pre032 | 10.10.2019 | 22 | m | Pre083 | 22.10.2019 | 18 | m | Pre134 | 12.11.2019 | 37 | w |
| Pre033 | 10.10.2019 | 29 | m | Pre084 | 22.10.2019 | 18 | w | Pre135 | 12.11.2019 | 55 | w |
| Pre034 | 11.10.2019 | 42 | m | Pre085 | 22.10.2019 | 52 | m | Pre136 | 12.11.2019 | 59 | m |
| Pre035 | 11.10.2019 | 27 | m | Pre086 | 22.10.2019 | 52 | m | Pre137 | 14.11.2019 | 25 | m |
| Pre036 | 11.10.2019 | 27 | m | Pre087 | 22.10.2019 | 37 | w | Pre138 | 14.11.2019 | 23 | w |
| Pre037 | 11.10.2019 | 55 | m | Pre088 | 22.10.2019 | 20 | m | Pre139 | 14.11.2019 | 20 | m |
| Pre038 | 11.10.2019 | 50 | m | Pre089 | 23.10.2019 | 20 | m | Pre140 | 14.11.2019 | 27 | w |
| Pre039 | 11.10.2019 | 55 | m | Pre090 | 23.10.2019 | 27 | w | Pre141 | 14.11.2019 | 20 | m |
| Pre040 | 11.10.2019 | 31 | m | Pre091 | 23.10.2019 | 19 | m | Pre142 | 14.11.2019 | 42 | m |
| Pre041 | 11.10.2019 | 25 | w | Pre092 | 23.10.2019 | 25 | w | Pre143 | 14.11.2019 | 35 | w |
| Pre042 | 11.10.2019 | 22 | w | Pre093 | 23.10.2019 | 27 | m | Pre144 | 14.11.2019 | 42 | w |
| Pre043 | 11.10.2019 | 32 | m | Pre094 | 23.10.2019 | 39 | m | Pre145 | 15.11.2019 | 40 | m |
| Pre044 | 11.10.2019 | 45 | m | Pre095 | 23.10.2019 | 25 | w | Pre146 | 15.11.2019 | 59 | w |
| Pre045 | 11.10.2019 | 28 | w | Pre096 | 23.10.2019 | 20 | w | Pre147 | 15.11.2019 | 52 | w |
| Pre046 | 15.10.2019 | 19 | w | Pre097 | 23.10.2019 | 58 | m | Pre148 | 15.11.2019 | 41 | m |
| Pre047 | 15.10.2019 | 25 | w | Pre098 | 24.10.2019 | 18 | w | Pre149 | 15.11.2019 | 24 | w |
| Pre048 | 15.10.2019 | 20 | m | Pre099 | 24.10.2019 | 22 | w | Pre150 | 15.11.2019 | 23 | m |
| Pre049 | 15.10.2019 | 44 | m | Pre100 | 24.10.2019 | 19 | m | EV005 | 16.12.2016 | 44 | m |
| Pre050 | 15.10.2019 | 28 | w | Pre101 | 24.10.2019 | 20 | w | EV006 | 20.01.2017 | 45 | m |
| Pre051 | 15.10.2019 | 23 | m | Pre102 | 24.10.2019 | 19 | w | | | | |

**Supplementary table 2: Sequence information of pre-pandemic and original heavy and light chains**

| NO | SARS-COV-2- REACTIVE AB | PRE-PANDEMIC CHAIN | NAME IN FIGURE | PRE-PANDEMIC AB NAME | SARS-COV-2- REACTIVE AB V AA | PRE-PANDEMIC CHAIN V AA | V DISTANCE AA | SARS-COV-2- REACTIVE AB CDR3 AA | PRE-PANDEMIC CHAIN CDR3 AA | CDR3 DISTANCE AA |
|---|---|---|---|---|---|---|---|---|---|---|
| 1 | CnC2t1p1_B4 | HC | HC1 | CnC2t1p1_B4_pc-1 | QVQLVQSGAEVKKPGASVKVSCKASGYTFTSYGISWVRQAPGQGLEWMGWISAYNGNTNYAQKLQGRVTMTTDTSTSTAYMELRSLRSDDTAVYYC | QVQLVQSGAEVKKPGASVKVSCKASGYTFTSYGISWVRQAPGQGLEWMGWISAYNGNTNYAQKLQGRVTMTTDTSTSTAYMELRSLRSDDTAVYYC | 0 | ARDGELLGWFDP | ARDSPLGWFDP | 3 |
| 2 | CnC2t1p1_B4 | HC | HC2 | CnC2t1p1_B4_pc-2 | QVQLVQSGAEVKKPGASVKVSCKASGYTFTSYGISWVRQAPGQGLEWMGWISAYNGNTNYAQKLQGRVTMTTDTSTSTAYMELRSLRSDDTAVYYC | QVQLVQSGAEVKKPGASVKVSCKASGYTFTSYGISWVRQAPGQGLEWMGWISAYNGNTNYAQKLQGRVTMTTDTSTSTAYMELRSLRSDDTAVYYC | 0 | ARDGELLGWFDP | ARGEWELGWFDP | 3 |
| 3 | HbnC3t1p1_G4 | HC | HC1 | HbnC3t1p1_G4_pc-2 | EVQLVESGGGLVQPGGSLRLSCAASGFTVSSNYMSWVRQAPGKGLEWVSVIYSGGSTFYADSVKGRFTISRDNSKNTLYLQMNSLRAEDTAVYYC | EVQLVESGGGLVQPGGSLRLSCAASGFTVSSNYMSWVRQAPGKGLEWVSVIYSGGSTYYADSVKGRFTISRDNSKNTLYLQMNSLRAEDTAVYYC | 1 | ARDFGDFFFDY | ARDNGDYYFDY | 3 |
| 4 | HbnC3t1p2_B10 | HC | HC1 | HbnC3t1p2_B10_pc-1 | EVQLVESGGGLVQPGGSLRLSCAASGFIVSSNYMSWVRQAPGKGLEWVSVIYSGGSTFYADSVKGRFTISRDNSKNTLYLQMNSLRAEDTAVYYC | EVQLVESGGGLVQPGGSLRLSCAASGFTVSSNYMSWVRQAPGKGLEWVSVIYSGGSTYYADSVKGRFTISRDNSKNTLYLQMNSLRAEDTAVYYC | 2 | ARDYGDYFFDY | ARDNGDYYFDY | 2 |
| 5 | HbnC3t1p2_B10 | HC | HC2 | HbnC3t1p2_B10_pc-2 | EVQLVESGGGLVQPGGSLRLSCAASGFIVSSNYMSWVRQAPGKGLEWVSVIYSGGSTFYADSVKGRFTISRDNSKNTLYLQMNSLRAEDTAVYYC | EVQLVESGGGLVQPGGSLRLSCAASGFTVSSNYMSWVRQAPGKGLEWVSVIYSGGSTYYADSVKGRFTISRDNSKNTLYLQMNSLRAEDTAVYYC | 2 | ARDYGDYFFDY | ARAYGDYFDY | 2 |
| 6 | HbnC3t1p2_B10 | HC | HC3 | HbnC3t1p2_B10_pc-3 | EVQLVESGGGLVQPGGSLRLSCAASGFIVSSNYMSWVRQAPGKGLEWVSVIYSGGSTFYADSVKGRFTISRDNSKNTLYLQMNSLRAEDTAVYYC | EVQLVESGGGLVQPGGSLRLSCAASGFTVSSNYMSWVRQAPGKGLEWVSVIYSGGSTYYADSVKGRFTISRDNSKNTLYLQMNSLRAEDTAVYYC | 2 | ARDYGDYFFDY | ARDPGGYYFDY | 3 |
| 7 | HbnC3t1p2_B10 | HC | HC4 | HbnC3t1p2_B10_pc-4 | EVQLVESGGGLVQPGGSLRLSCAASGFIVSSNYMSWVRQAPGKGLEWVSVIYSGGSTFYADSVKGRFTISRDNSKNTLYLQMNSLRAEDTAVYYC | EVQLVESGGGLVQPGGSLRLSCAASGFTVSSNYMSWVRQAPGKGLEWVSVIYSGGSTYYADSVKGRFTISRDNSKNTLYLQMNSLRAEDTAVYYC | 2 | ARDYGDYFFDY | ARDYNGDYSFDY | 2 |
| 8 | HbnC3t1p2_B10 | HC | HC5 | HbnC3t1p2_B10_pc-5 | EVQLVESGGGLVQPGGSLRLSCAASGFIVSSNYMSWVRQAPGKGLEWVSVIYSGGSTFYADSVKGRFTISRDNSKNTLYLQMNSLRAEDTAVYYC | EVQLVESGGGLVQPGGSLRLSCAASGFTVSSNYMSWVRQAPGKGLEWVSVIYSGGSTYYADSVKGRFTISRDNSKNTLYLQMNSLRAEDTAVYYC | 2 | ARDYGDYFFDY | ARGYGDYPLDY | 3 |
| 9 | HbnC4t1p1_A1 | HC | HC1 | HbnC4t1p1_A1_pc-1 | EVQLLESGGGLVQPGGSLRLSCAASGFTFSSYAMSWVRQAPGKGLEWVSAISGSGGSTYYADSVKGRFTISRDNSKNTLYLQMNSLRAEDTAVYYC | EVQLLESGGGLVQPGGSLRLSCAASGFTFSSYAMSWVRQAPGKGLEWVSAISGSGGSTYYADSVKGRFTISRDNSKNTLYLQMNSLRAEDTAVYYC | 0 | AKALAAAGYWAFDY | AKAIAAAGYYFDY | 3 |
| 10 | HbnC4t1p1_C11 | HC | HC1 | HbnC4t1p1_C11_pc-1 | EVQLLESGGGLVQPGGSLRLSCAASGFTFSSYAMSWVRQAPGKGLEWVSAISGSGGSTYYADSVKGRFTISRDNSKNTLYLQMNSLRAEDTAVYYC | EVQLLESGGGLVQPGGSLRLSCAASGFTFSSYAMSWVRQAPGKGLEWVSAISGSGGSTYYADSVKGRFTISRDNSKNTLYLQMNSLRAEDTAVYYC | 0 | AKALAAAGYWAFDY | AKAIAAAGYYFDY | 3 |
| 11 | MnC2t1p1_C12 | HC | HC1 | MnC2t1p1_C12_pc-1 | QVQLQESGPGLVKPSQTLSLTCTVSGGSISSSGYFWSWIRQHPGKGLEWIGYIYYSGSTYYNPSLKSRVTISVDTSKNQFSLKLSSLTAADTAVYYC | QVQLQESGPGLVKPSQTLSLTCTVSGGSISSGGYYWSWIRQHPGKGLEWIGYIYYSGSTYYNPSLKSRVTISVDTSKNQFSLKLSSVTAADTAVYYC | 3 | ARVGSGRAFDI | ARGGSRDAFDI | 3 |
| 12 | MnC2t1p1_C12 | HC | HC2 | MnC2t1p1_C12_pc-2 | QVQLQESGPGLVKPSQTLSLTCTVSGGSISSSGYFWSWIRQHPGKGLEWIGYIYYSGSTYYNPSLKSRVTISVDTSKNQFSLKLSSLTAADTAVYYC | QVQLQESGPGLVKPSQTLSLTCTVSGGSISSGGYYWSWIRQHPGKGLEWIGYIYYSGSTYYNPSLKSRVTISVDTSKNQFSLKLSSVTAADTAVYYC | 3 | ARVGSGRAFDI | ARVGSPHDAFDI | 3 |
| 13 | MnC2t1p1_C12 | HC | HC3 | MnC2t1p1_C12_pc-3 | QVQLQESGPGLVKPSQTLSLTCTVSGGSISSSGYFWSWIRQHPGKGLEWIGYIYYSGSTYYNPSLKSRVTISVDTSKNQFSLKLSSLTAADTAVYYC | QVQLQESGPGLVKPSQTLSLTCTVSGGSISSGGYYWSWIRQHPGKGLEWIGYIYYSGSTYYNPSLKSRVTISVDTSKNQFSLKLSSVTAADTAVYYC | 3 | ARVGSGRAFDI | ARVGSGYDAFDI | 2 |
| 14 | MnC2t1p1_C12 | HC | HC4 | MnC2t1p1_C12_pc-4 | QVQLQESGPGLVKPSQTLSLTCTVSGGSISSSGYFWSWIRQHPGKGLEWIGYIYYSGSTYYNPSLKSRVTISVDTSKNQFSLKLSSLTAADTAVYYC | QVQLQESGPGLVKPSQTLSLTCTVSGGSISSGGYYWSWIRQHPGKGLEWIGYIYYSGSTYYNPSLKSRVTISVDTSKNQFSLKLSSVTAADTAVYYC | 3 | ARVGSGRAFDI | ARVGSKAFDI | 2 |
| 15 | MnC2t1p1_C12 | HC | HC5 | MnC2t1p1_C12_pc-5 | QVQLQESGPGLVKPSQTLSLTCTVSGGSISSSGYFWSWIRQHPGKGLEWIGYIYYSGSTYYNPSLKSRVTISVDTSKNQFSLKLSSLTAADTAVYYC | QVQLQESGPGLVKPSQTLSLTCTVSGGSISSGGYYWSWIRQHPGKGLEWIGYIYYSGSTYYNPSLKSRVTISVDTSKNQFSLKLSSVTAADTAVYYC | 3 | ARVGSGRAFDI | ARAGSFDAFDI | 3 |
| 16 | MnC2t1p1_C12 | HC | HC6 | MnC2t1p1_C12_pc-6 | QVQLQESGPGLVKPSQTLSLTCTVSGGSISSSGYFWSWIRQHPGKGLEWIGYIYYSGSTYYNPSLKSRVTISVDTSKNQFSLKLSSLTAADTAVYYC | QVQLQESGPGLVKPSQTLSLTCTVSGGSISSGGYYWSWIRQHPGKGLEWIGYIYYSGSTYYNPSLKSRVTISVDTSKNQFSLKLSSVTAADTAVYYC | 3 | ARVGSGRAFDI | ARVGSGRAFDI | 0 |
| 17 | MnC2t1p1_C12 | HC | HC7 | MnC2t1p1_C12_pc-7 | QVQLQESGPGLVKPSQTLSLTCTVSGGSISSSGYFWSWIRQHPGKGLEWIGYIYYSGSTYYNPSLKSRVTISVDTSKNQFSLKLSSLTAADTAVYYC | QVQLQESGPGLVKPSQTLSLTCTVSGGSISSGGYYWSWIRQHPGKGLEWIGYIYYSGSTYYNPSLKSRVTISVDTSKNQFSLKLSSVTAADTAVYYC | 3 | ARVGSGRAFDI | ARVGLNAFDI | 3 |
| 18 | MnC2t1p1_C12 | HC | HC8 | MnC2t1p1_C12_pc-8 | QVQLQESGPGLVKPSQTLSLTCTVSGGSISSSGYFWSWIRQHPGKGLEWIGYIYYSGSTYYNPSLKSRVTISVDTSKNQFSLKLSSLTAADTAVYYC | QVQLQESGPGLVKPSQTLSLTCTVSGGSISSGGYYWSWIRQHPGKGLEWIGYIYYSGSTYYNPSLKSRVTISVDTSKNQFSLKLSSVTAADTAVYYC | 3 | ARVGSGRAFDI | ARAGSRGAFDI | 3 |
| 19 | MnC2t1p1_C12 | HC | HC9 | MnC2t1p1_C12_pc-9 | QVQLQESGPGLVKPSQTLSLTCTVSGGSISSSGYFWSWIRQHPGKGLEWIGYIYYSGSTYYNPSLKSRVTISVDTSKNQFSLKLSSLTAADTAVYYC | QVQLQESGPGLVKPSQTLSLTCTVSGGSISSGGYYWSWIRQHPGKGLEWIGYIYYSGSTYYNPSLKSRVTISVDTSKNQFSLKLSSVTAADTAVYYC | 3 | ARVGSGRAFDI | ARVPLGAFDI | 3 |
| 20 | MnC2t1p1_C12 | HC | HC10 | MnC2t1p1_C12_pc-11 | QVQLQESGPGLVKPSQTLSLTCTVSGGSISSSGYFWSWIRQHPGKGLEWIGYIYYSGSTYYNPSLKSRVTISVDTSKNQFSLKLSSLTAADTAVYYC | QVQLQESGPGLVKPSQTLSLTCTVSGGSISSGGYYWSWIRQHPGKGLEWIGYIYYSGSTYYNPSLKSRVTISVDTSKNQFSLKLSSVTAADTAVYYC | 3 | ARVGSGRAFDI | ARVGIFDAFDI | 3 |
| 21 | MnC2t1p1_C12 | HC | HC11 | MnC2t1p1_C12_pc-12 | QVQLQESGPGLVKPSQTLSLTCTVSGGSISSSGYFWSWIRQHPGKGLEWIGYIYYSGSTYYNPSLKSRVTISVDTSKNQFSLKLSSLTAADTAVYYC | QVQLQESGPGLVKPSQTLSLTCTVSGGSISSGGYYWSWIRQHPGKGLEWIGYIYYSGSTYYNPSLKSRVTISVDTSKNQFSLKLSSVTAADTAVYYC | 3 | ARVGSGRAFDI | AREGWGAFDI | 3 |
| 22 | MnC2t1p1_C12 | HC | HC12 | MnC2t1p1_C12_pc-13 | QVQLQESGPGLVKPSQTLSLTCTVSGGSISSSGYFWSWIRQHPGKGLEWIGYIYYSGSTYYNPSLKSRVTISVDTSKNQFSLKLSSLTAADTAVYYC | QVQLQESGPGLVKPSQTLSLTCTVSGGSISSGGYYWSWIRQHPGKGLEWIGYIYYSGSTYYNPSLKSRVTISVDTSKNQFSLKLSSVTAADTAVYYC | 3 | ARVGSGRAFDI | ARVPTGARAFDI | 3 |
| 23 | MnC2t1p1_C12 | HC | HC13 | MnC2t1p1_C12_pc-14 | QVQLQESGPGLVKPSQTLSLTCTVSGGSISSSGYFWSWIRQHPGKGLEWIGYIYYSGSTYYNPSLKSRVTISVDTSKNQFSLKLSSLTAADTAVYYC | QVQLQESGPGLVKPSQTLSLTCTVSGGSISSGGYYWSWIRQHPGKGLEWIGYIYYSGSTYYNPSLKSRVTISVDTSKNQFSLKLSSVTAADTAVYYC | 3 | ARVGSGRAFDI | ARVGGYRAFDI | 2 |
| 24 | CnC2t1p1_B4 | LC | LC1 | CnC2t1p1_B4_pc-1 | QSALTQPASVSGSPGQSITISCTGTSSDVGSYNLVSWYQQHPGKAPKLMIYEGSKRPSGVSNRFSGSKSGNTASLTISGLQAEDEADYYC | QSALTQPASVSGSPGQSITISCTGTSSDVGSYNLVSWYQQHPGKAPKLMIYEGSKRPSGVSNRFSGSKSGNTASLTISGLQAEDEADYYCC | 1 | CSYAGSSTWV | CSYAGSSTWV | 0 |
| 25 | HbnC3t1p1_G4 | KC | LC1 | HbnC3t1p1_G4_pc-2 | EIVLTQSPGTLSLSPGERATLSCRASQSVSSYLAWYQQKPGQAPRLLIYGVSSRATGIPDRFSGSGSGTDFTLTISRLEPEDFAVYYC | EIVLTQSPGTLSLSPGERATLSCRASQSVSSSYLAWYQQKPGQAPRLLIYGASSRATGIPDRFSGSGSGTDFTLTISRLEPEDFAVYYC | 2 | QQYGSSPRT | QQYGSSPRT | 0 |
| 26 | HbnC3t1p2_B10 | KC | LC1 | HbnC3t1p2_B10_pc-1 | EIVLTQSPGTLSLSPGERATLSCRASQSVSSYLAWYQQKPGQAPRLLISGASSRAAGIPDRFSGSGSGTDFTLTINRLEPEDFAVYYC | EIVLTQSPGTLSLSPGERATLSCRASQSVSSSYLAWYQQKPGQAPRLLIYGASSRATGIPDRFSGSGSGTDFTLTISRLEPEDFAVYYC | 4 | QQYGSSPRT | QQYGSSPRT | 0 |
| 27 | HbnC4t1p1_A1 | KC | LC1 | HbnC4t1p1_A1_pc-1 | DIQMTQSPSSLSASVGDRVTITCQASQDISNYLNWYQQKPGKAPKLLIYDASNLETGVPSRFSGSGSGTDFTFTISSLQPEDIATYYC | DIQMTQSPSSLSASVGDRVTITCQASQDISNYLNWYQQKPGKAPKLLIYDASNLETGVPSRFSGSGSGTDFTFTISSLQPEDIATYYC | 0 | QQYDNLRLT | QQYDNLVLT | 1 |
| 28 | HbnC4t1p1_C11 | KC | LC1 | HbnC4t1p1_C11_pc-1 | DIQMTQSPSSLSASVGDRVTITCQASQDISNYLNWYQQKPGKAPKLLIYDASNLETGVPSRFSGSGSGTDFTFTISSLQPEDIATYYC | DIQMTQSPSSLSASVGDRVTITCQASQDISNYLNWYQQKPGKAPKLLIYDASNLETGVPSRFSGSGSGTDFTFTISSLQPEDIATYYC | 0 | QQYDNLRLT | QQYDNLVLT | 1 |
| 29 | MnC2t1p1_C12 | KC | LC1 | MnC2t1p1_C12_pc-1 | EIVLTQSPVTLSLSPGERATLSCRASQSLSDSYLAWYQQKPGQAPRLLIYNASSRATGIPDRFSGSGSGTDFTLTISRLAPEDFAVYYC | EIVLTQSPGTLSLSPGERATLSCRASQSVSSSYLAWYQQKPGQAPRLLIYGASSRATGIPDRFSGSGSGTDFTLTISRLEPEDFAVYYC | 5 | QYYGSSL | QQYGSSL | 1 |
| 30 | MnC2t1p1_C12 | KC | LC2 | MnC2t1p1_C12_pc-7 | EIVLTQSPVTLSLSPGERATLSCRASQSLSDSYLAWYQQKPGQAPRLLIYNASSRATGIPDRFSGSGSGTDFTLTISRLAPEDFAVYYC | EIVLTQSPGTLSLSPGERATLSCRASQSVSSSYLAWYQQKPGQAPRLLIYGASSRATGIPDRFSGSGSGTDFTLTISRLEPEDFAVYYC | 5 | QYYGSSL | QQYGSST | 2 |
| 31 | MnC2t1p1_C12 | KC | LC3 | MnC2t1p1_C12_pc-12 | EIVLTQSPVTLSLSPGERATLSCRASQSLSDSYLAWYQQKPGQAPRLLIYNASSRATGIPDRFSGSGSGTDFTLTISRLAPEDFAVYYC | EIVLTQSPGTLSLSPGERATLSCRASQSVSSSYLAWYQQKPGQAPRLLIYGASSRATGIPDRFSGSGSGTDFTLTISRLEPEDFAVYYC | 5 | QYYGSSL | QQYGSSP | 2 |
